# Machine Learning–Guided Differentiation Therapy Targets Cancer Stem Cells in Colorectal Cancers

**DOI:** 10.1101/2023.09.13.557628

**Authors:** Saptarshi Sinha, Joshua Alcantara, Kevin Perry, Vanessa Castillo, Annelies Ondersma, Satarupa Banerjee, Ella McLaren, Celia R. Espinoza, Sahar Taheri, Eleadah Vidales, Courtney Tindle, Adel Adel, Siamak Amirfakhri, Joseph R. Sawires, Jerry Yang, Michael Bouvet, Pradipta Ghosh

## Abstract

Despite advances in artificial intelligence (AI) within cancer research, its application toward realizing differentiation therapy in solid tumors remains limited. Using colorectal cancer (CRC) as a model, we developed a machine learning (ML) framework, ***CANDiT*** (*Cancer Associated Nodes for Differentiation Targeting*), to selectively induce differentiation and death of cancer stem cells (CSCs)—a key obstacle to durable response. Centering on one node, *CDX2*, a master differentiation factor lost in high-risk, poorly differentiated CRCs, we built a transcriptomic network to identify therapeutic strategies for CDX2 restoration. Network-based prioritization identified *PRKAB1*, a stress polarity sensor, as a top target. A clinical-grade PRKAB1 agonist reprogrammed transcriptional networks, induced crypt differentiation, and selectively eliminated CDX2-low CSCs in CRC cell lines, xenografts and patient-derived organoids (PDOs). Multivariate analyses in PDOs revealed a strong therapeutic index, linking efficacy (IC₅₀) to the biomarker-defined CDX2-low state. A 50-gene response signature—derived from an integrated analyses of all three models and trained across multiple datasets—revealed that CDX2 restoration therapy may translate into a ∼50% reduction in recurrence and mortality risk. Mechanistically, treatment activated a differentiation-associated stress polarity signaling axis while dismantling Wnt and YAP-driven stemness programs essential to CSC survival. Thus, *CANDiT* offers a scalable path to CSC–directed therapy in solid tumors by translating transcriptomic vulnerabilities into precision treatments.

**Graphic Abstract:** 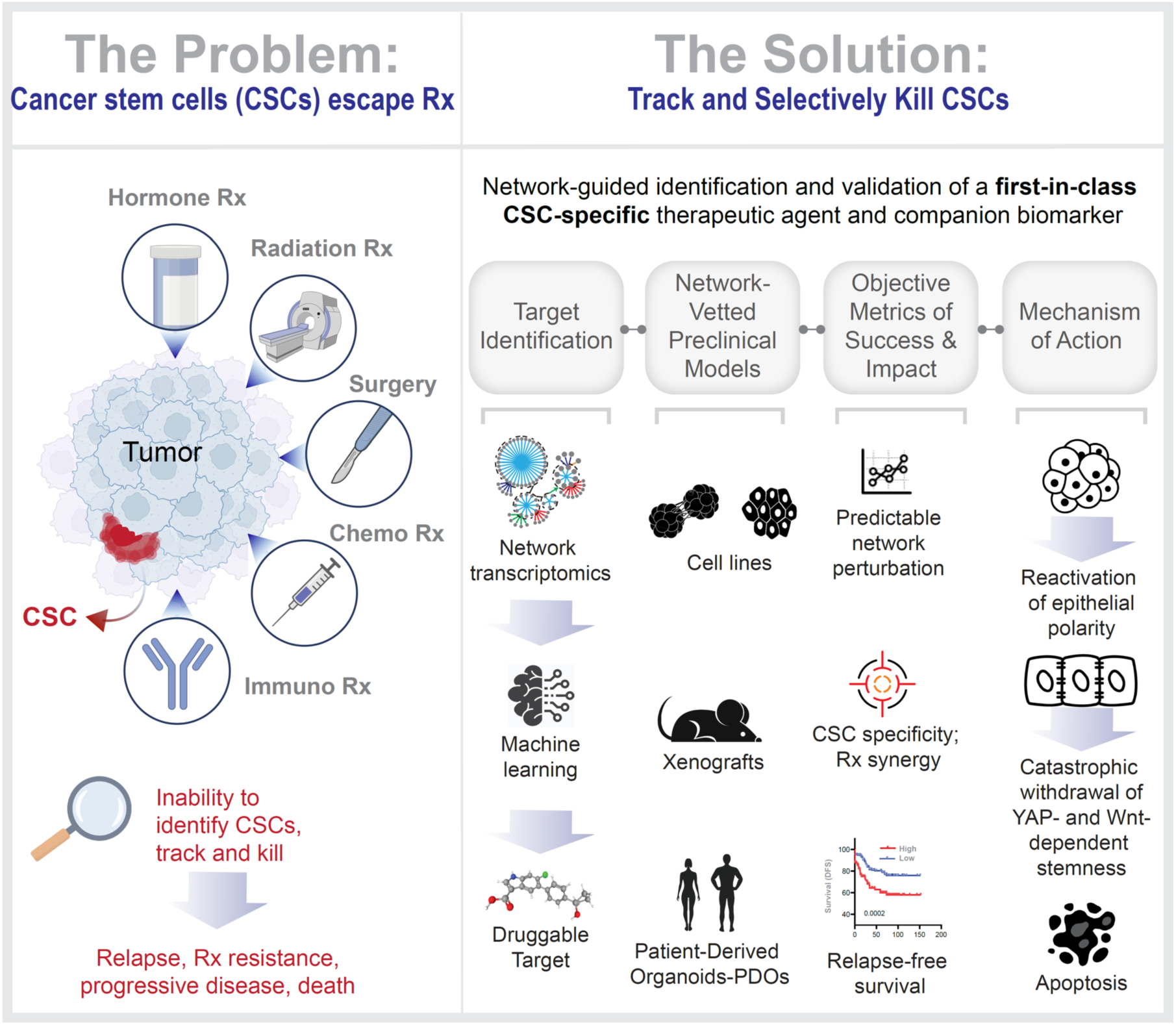

**One sentence summary:** In this work, Sinha et al. introduce a machine learning–guided framework to identify and target transcriptomic vulnerabilities in colorectal cancer, demonstrating that differentiation therapy selectively eliminates cancer stem cells and reduces recurrence risk.

**Highlights:**

- An ML framework (*CANDiT*) identifies target for differentiation therapy for CRCs
- Therapy induces crypt differentiation and CSC-specific cytotoxicity
- CDX2-low state predicts therapeutic response; restoration improves prognosis
- Therapy dismantles stemness via reactivation of stress polarity signaling

## Introduction

Poor differentiation, marked by elevated stemness, is a hallmark of cancers. Differentiation therapy, which targets cancer stem cells (CSCs) to induce maturation, has shown success in hematologic malignancies— most notably with all-trans-retinoic acid (ATRA) in acute promyelocytic leukemia (APML)^1–10^. While other agents have also shown promise in leukemias^11,12^, differentiation therapy has not yet translated to carcinomas. A major obstacle is the profound intra- and inter-tumoral heterogeneity that obscures identification of CSCs^13^.

To address this, computational methods have been developed to interrogate transcriptomic data, construct gene networks, and identify therapeutic targets^14–19^. Traditional symmetric frameworks—such as correlation^20–25^, mutual information^17^, linear regression^26^, dimension reduction^27^, and clustering^28,29^ —often fail in the face of real-world biological complexity. In contrast, Boolean implication-based network transcriptomics^30,31^ uses asymmetric, invariant gene relationships to construct directed networks that map evolving cellular states. This approach has identified translationally relevant states and targets across tissues and contexts^32–45^ with high degrees of precision, including a first-in-class therapeutic to protect gut barrier function in inflammatory bowel diseases (IBD)^30^.

Here we applied this method to identify targets that restore expression of CDX2, a caudal-related homeobox transcription factor and tumor suppressor^46–49^. In 2016, an unbiased search (using the same Boolean logic approach^50,51^) identified CDX2 as the top marker of colonic epithelial differentiation^52^. Its expression inversely correlates with “activated leukocyte cell adhesion molecule” (*ALCAM*/CD166), a stem cell marker that is present at the crypt base^53,54^ and on highly tumorigenic human CRC cells^55^. CDX2 loss— seen in ∼9% of CRCs—correlates with poor differentiation, worse prognosis, and enhanced chemotherapy benefit even in stage II disease^52^. Since then, 32 independent studies involving >13,000 patients have confirmed CDX2-low CRCs are associated with worse overall (OS) and disease-free (DFS) survival, independent of stage, mismatch repair (MMR) status, or ethnicity^56–70^ [**Figure 1**-*Step 1*]. One study integrated IHC with quantitative mass spectrometry^71^. Absence of CDX2 in tumors is associated with several adverse prognostic features, such as poor differentiation, advanced stage, vascular invasion, right-sided location, CpG island methylator phenotype (CIMP), and *BRAF* mutation^72–74^. Conversely, CDX2 presence lowers recurrence and mortality risk by ∼50% and 52%, respectively^69^ —with even greater benefit (∼70%) in stage II–III CRCs. Collectively, these findings establish the restoration of CDX2 as a clinically meaningful and high-priority therapeutic goal [see **Figure 1**-*Step 1*]. While its biological importance is widely recognized, no pharmacologic approach has yet succeeded in reliably inducing *CDX2*. Because *CDX2* alone is insufficient to fully activate the intestinal lineage program^75^ —and that its precise regulation is critical to preserve mucosal architecture—exogenous expression approaches are impractical.

**Figure 1.**
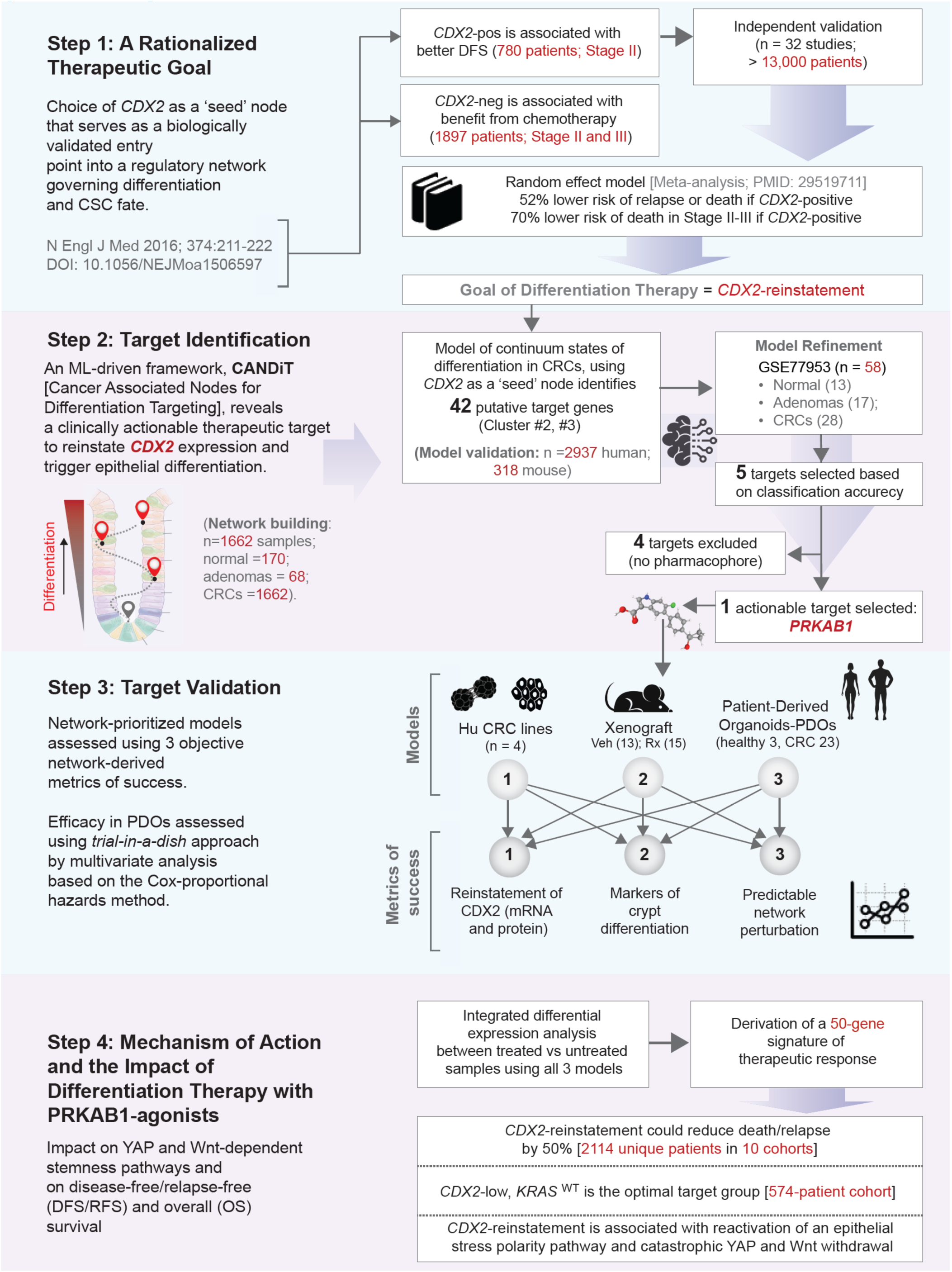
Study Design: Logical Network Perturbation to Reinvigorate CDX2 Expression as a Therapeutic Strategy in CRCs. **Step 1**: Flowchart outlines the chronological accumulation of evidence supporting *CDX2*’s biological role as a master regulator of intestinal epithelial differentiation and its consistent association with poor outcomes^76^. **Step 2**. Key steps in the network-based identification of PRKAB1 as a potent, actionable target, whose agonism is predicted to restore CDX2 expression. See **Supplemental Information 1** for datasets used in model training and validation, and **Supplemental Information 2** for gene clusters co-induced with *CDX2* upon PRKAB1 activation. **Step 3.** Three experimental models and objective criteria used to validate PRKAB1 agonism as a strategy for CDX2 reinstatement and induction of cellular differentiation. **Step 4**. Computationally driven insights into underlying mechanism of action and therapeutic impact. CDX2-reinstatement may halve the risk of death/relapse and is associated with a catastrophic collapse of signals that sustain cancer stemness---programs to which CSCs, but not other cancer cells are addicted--and with the reactivation of an epithelial stress polarity pathway that is selectively lost in cancers.

Here, we used network-transcriptomic analysis^50,51^ to identify therapeutic targets that upregulate CDX2, triggering a network-wide, domino-like cascade that drives differentiation in CRC. We selected CDX2 as a biologically validated master regulator whose reactivation enables reprograms CSC fate^76^, and bridges cellular differentiation states to clinical outcomes, linking cancer stem cell behavior to patient prognosis.

## Results

### A network-rationalized approach, CANDiT

We applied a machine learning–based computational platform, the Boolean Network Explorer (*BoNE*)^30^, and *CDX2* as a ‘seed’ gene (i.e., node) to model progressive gene regulatory events during colonic differentiation (**Figure 1**-*Step 2;* **Figure S1A-B**). The resulting framework—*CANDiT* (*Cancer-Associated Nodes for Differentiation Targeting*)—prioritizes actionable differentiation nodes in cancer networks to enable therapeutic reprogramming. The model was trained initially on a maximally heterogeneous dataset comprising 1,662 colorectal cancers (CRCs), 68 adenomas, and 170 normal tissues (**Figure S1C-D**). The model was refined subsequently using statistical learning on another independent dataset (<u>GSE77953</u>). Classification accuracy identified five statistically top-ranked target genes from *CDX2*-proximal clusters (#2 and #3; **Figure S1D**) that are predicted to upregulate *CDX2* when activated (**Figure 1**-*Step 2;* **Figure S1E**). These predictions were validated in 2,937 additional human colon samples (**Figure S1F**) and 318 mouse samples (**Figure S1G**), confirming cross-species and cross-cohort robustness.

To identify potent, selective modulators of the five targets, we queried the Protein Data Bank [PDB; https://www.rcsb.org] for structure-resolved agonists. Selective ligands were identified only for PRKAB1 (25 molecules/structures); none were found for the other targets. Cross-referencing these hits in ClinicalTrials.gov yielded PF-06409577 (PF), a clinical-grade PRKAB1 agonist. Although not yet approved for a clinical indication, PF has been deemed safe in a randomized, double-blind, placebo-controlled Phase I trial (NCT02286882^77^). PF was recently shown to specifically upregulate PRKAB1 at both transcript and protein levels in the colonic epithelium ex vivo and in mouse colon in vivo^30^.

The *CANDiT*-inferred network predicted that agonizing PRKAB1, the protein product of *PRKAB1* (Protein Kinase AMP-activated Non-Catalytic Subunit Beta-1), would: (i) upregulate *PRKAB1* itself and co-clustered genes (via “equivalent” relationships); (ii) induce *CDX2* and its proximal clusters (#2 and #3) via “hi⇒hi” links; and (iii) repress stemness-associated genes (e.g., *CCDC88A*, cluster #9) via “hi⇒lo” links (**Figure S1D**). This regulatory logic holds consistently across diverse human and mouse CRC datasets, including early-onset CRCs (**Figure S2A-D**). Further supporting its selection, *PRKAB1*—but not its paralog *PRKAB2*—is highly expressed in the gastrointestinal tract^30^ and PRKAB1 agonists promote epithelial polarity in the gut^78–81^.

For validation studies *in cellulo* (CRC cell lines), *in vivo* (murine xenografts) and *ex vivo* (a ‘living biobank’ of patient derived organoids; PDOs), we prioritized models with contrasting CDX2 expression (high vs. low, as determined based on a threshold that was determined using *StepMiner*^82^; see *Methods*): CDX2-low (target phenotype) and CDX2-high (negative control). In addition to conventional anti-tumor readouts, three pre-specified success metrics were used to evaluate therapeutic efficacy; all three models were required to meet all three endpoints (see **Figure 1**-*Step 3*). Previous attempts to induce CDX2 transiently^83^, failed to produce durable differentiation and showed inconsistent impact across these same metrics (**Figure S2E-H**). Finally, through an integrated transcriptomic approach validated in tumor tissues, we define the molecular mechanism of action and translational impact of successful *CDX2* reinstatement via PRKAB1 agonism (see **Figure 1**-*Step 4*).

### Target validation on CRC cell lines and xenograft models

We prioritized two poorly differentiated CRC cell lines—HCT116 and SW480—which exhibited the lowest CDX2 transcript levels among 26 sequenced CRC cell lines profiled by microarray (<u>GSE10843</u>)—as optimal models for hypothesis testing. Two well-differentiated, *CDX2*-high cell lines (DLD1, *KRAS*^mut^ and Caco2, *KRAS*^WT^) were selected as a negative control (**Figure 2A**). PF induced cell death within 48 hours in CDX2-low HCT116 and SW480 cells (IC_50_ ∼6-7 µM), but not in *CDX2*-high DLD1 or Caco2 cells (IC₅₀ >20 µM; **Figure 2B**; **Figure S3A-C**). In both *CDX2*-low cell lines, PF-induced cell death was preceded by dose-dependent induction of differentiation and suppression of stemness markers, at both transcript (**Figure 2C**) and protein (**Figure S3D-G**) levels. These effects coincided with a dose-dependent increase in apoptosis (**Figure S4A-H**). Importantly, PF failed to reinstate CDX2, alter differentiation or stemness markers (**Figure 2D**), or induce cell death (**Figure 2E**) in *PRKAB1*-depleted cells, confirming target specificity of its anti-cancer effects. In contrast, Metformin—a non-specific, indirect agonist of AMP-kinase (activating both PRKAB1- and PRKAB2-containing complexes)—neither induced CDX2 expression nor triggered cell death, even at high doses (**Figure 2B**; **Figure S4I-K**), underscoring the necessity of PRKAB1-specific activation for therapeutic efficacy.

**Figure 2.**
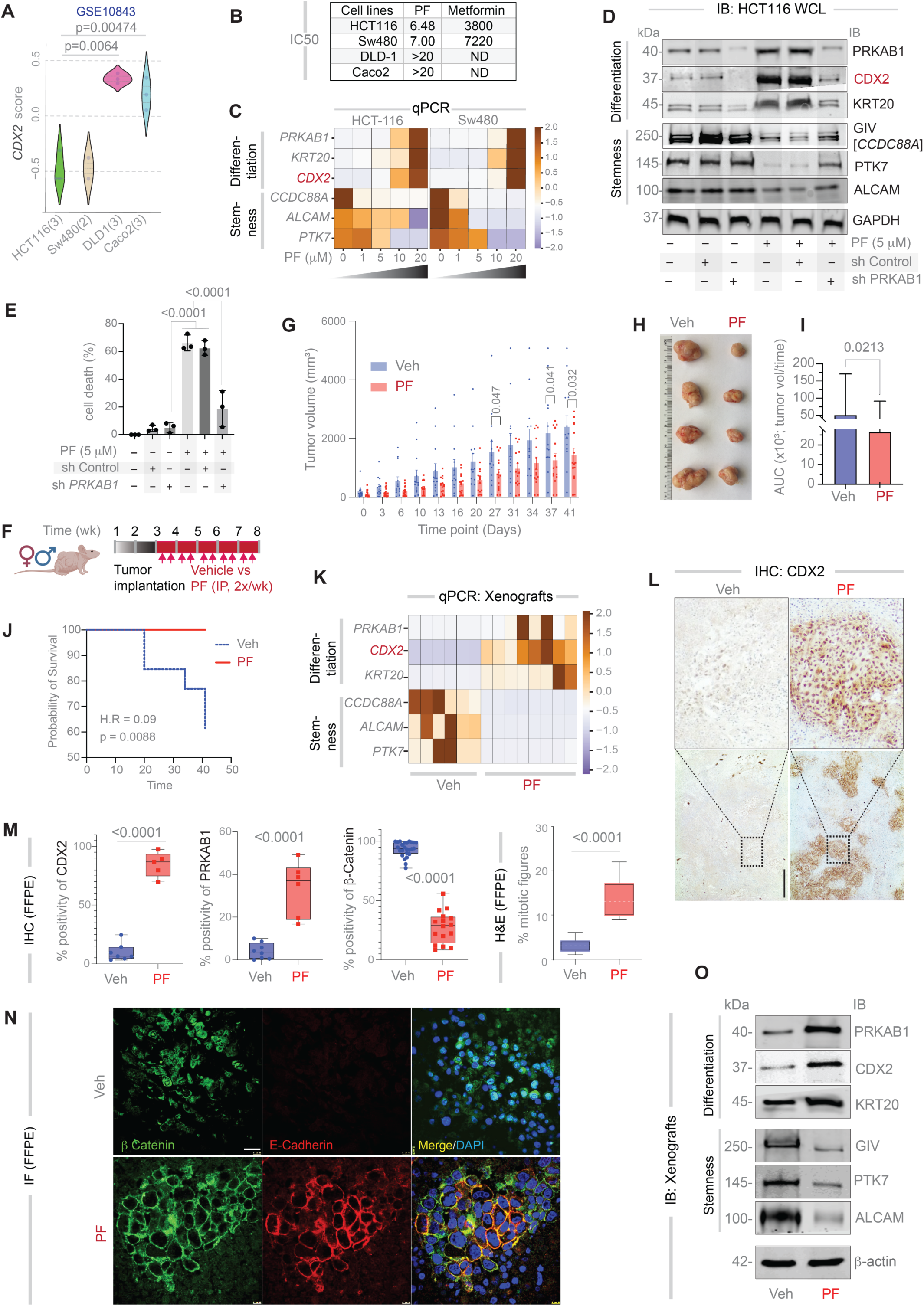
Target validation in CRC cell lines and xenograft models. **A.** Violin plot of normalized *CDX2* expression in CRC cell lines. *p*-values are calculated using Welch’s *t*-test. **B.** Table summarizing IC_50_ values (µM) for a specific (PF) or a non-specific (Metformin) PRKAB1-agonist in CRC cell lines, measured by MTT assays at 48 h. **C.** RT-qPCR analysis of differentiation and stemness markers in >95% viable cells (16 h post-treatment), displayed as a Z-score heatmap. **D-E**. Immunoblots (**D**) and MTT-based survival assays (**E**) showing the effects of PRKAB1 depletion (via shRNA in HCT116 cells) on the anti-cancer efficacy of PF. *P*-values via one-way ANOVA; error bars represent S.E.M. of three independent biological replicates. See **Figures S3** and **S4** for additional dose-response studies using PF (**S3**, **S4A–H**) and Metformin (**S4I–K**). **F.** Schematic of the xenograft experimental design and workflow. **G.** Tumor growth kinetics in vehicle-treated (Veh; *n* = 13) vs. PF-treated (*n* = 15) mice, measured twice weekly with calipers across three experimental cohorts. *P*-values by multiple paired *t*-tests with Benjamini, Krieger, and Yekutieli correction. **H.** Representative images of xenograft tumors at 8 weeks post-implantation. **I.** Bar plots comparing area under the curve (AUC) for tumor growth: Veh (95% CI: 31,788–70,772) vs. PF (95% CI: 16,949–36,404). *P*-value by unpaired two-tailed *t*-test; error bars indicate S.E.M. **J.** Kaplan–Meier analysis of time to IACUC-mandated endpoints (euthanasia due to death or discomfort). *P*-value was derived via log-rank test; H.R., hazard ratio. **K.** RT-qPCR analysis of differentiation and stemness markers in xenografts displayed as a heatmap of Z-score normalized values. **L.** Representative images of CDX2 immunostaining in FFPE xenografts. Scale bar = 100 µm. **M.** Quantification of CDX2, PRKAB1, and nuclear β-catenin (via IHC Profiler in ImageJ) and mitotic bodies (manually counted in H&E sections). p-values are calculated by unpaired 2-tailed *t*-test. See **Figure S5B-D** for images from representative fields. **N.** Representative fields from FFPE tumors co-stained for E-cadherin (red), β-catenin (green) and DAPI (nuclei; blue) are shown. Scale bar = 20 µm. See also **Figure S5K** for colocalization analysis using ImageJ. **O.** Immunoblots from viable tumor lysates. See **Figure S5F** for quantification.

In vivo, HCT116 xenografts implanted subcutaneously in nude mice exhibited a 68% reduction in tumor volume with PF treatment (AUC: vehicle = 51,280.23; PF = 26,676.77; **Figure 2G-I**) and a 62.5% increase in survival (**Figure 2J**; Mantel-Haenszel Hazard Ratio = 0.09 for reduction in death rates). PF-treated tumors developed glandular structures (**Figure S5A**), reinstated *CDX2* expression, and showed increased differentiation and reduced stemness markers by qPCR (**Figure 2K**). IHC confirmed nuclear localization of CDX2 and upregulation of PRKAB1 protein (IHC; **Figure 2L-M; Figure S5B**). Reflecting the slow division rate of colonic stem cells^84^, PF triggered a ∼3-fold increase in mitotic bodies (**Figure S5C**; **Figure 2M**). Additional signs of tumor differentiation included a reduction in nuclear β-catenin (**Figure S5D)** and a corresponding increase in junctional β-catenin (**Figure 2N**), which co-localized with E-cadherin (**Figure 2N; Figure S5E**).

Together, these findings demonstrate that CDX2-reinstatement therapy drives differentiation and apoptosis selectively in *CDX2*-low CRC models, both in vitro and in vivo, through PRKAB1-specific activation.

### Target validation in patient-derived organoids

We next evaluated the therapeutic efficacy of PF in PDOs. To evaluate potential toxicity, we first tested PF on healthy colon PDOs using eTOX Red staining, which selectively labels dead cells based on membrane permeability (**Figure 3A**-*left*). At 5 μM and 20 μM, PF, there was no significant increase in cell death compared to vehicle control. As expected, hydrogen peroxide (H₂O₂), a positive control, induced significant cytotoxicity relative to both PF-treated and control groups (**Figure 3A**-*right*; **Figure 3B**). We then tested PF across two prospective cohorts comprising 23 CRC PDOs and 3 healthy colon PDOs (see **Table** and *Methods*; **Figure 3C**). PF exhibited selective anti-cancer activity in CDX2-low CRC PDOs (as determined based on a threshold that was determined using *StepMiner*^82^ cutoff; see *Methods*), reflected by low IC₅₀ values. In contrast, CDX2-high CRC PDOs and all healthy colon PDOs were resistant (IC₅₀ > 40 µM; **Figure 3D**). Notably, IC₅₀ values positively correlated with expression of CDX2 (seed gene) and PRKAB1 (network-derived target), while showing an inverse correlation with CCDC88A, a network-linked stemness marker (**Figure S1D**; **Figure 3D**).

**Figure 3.**
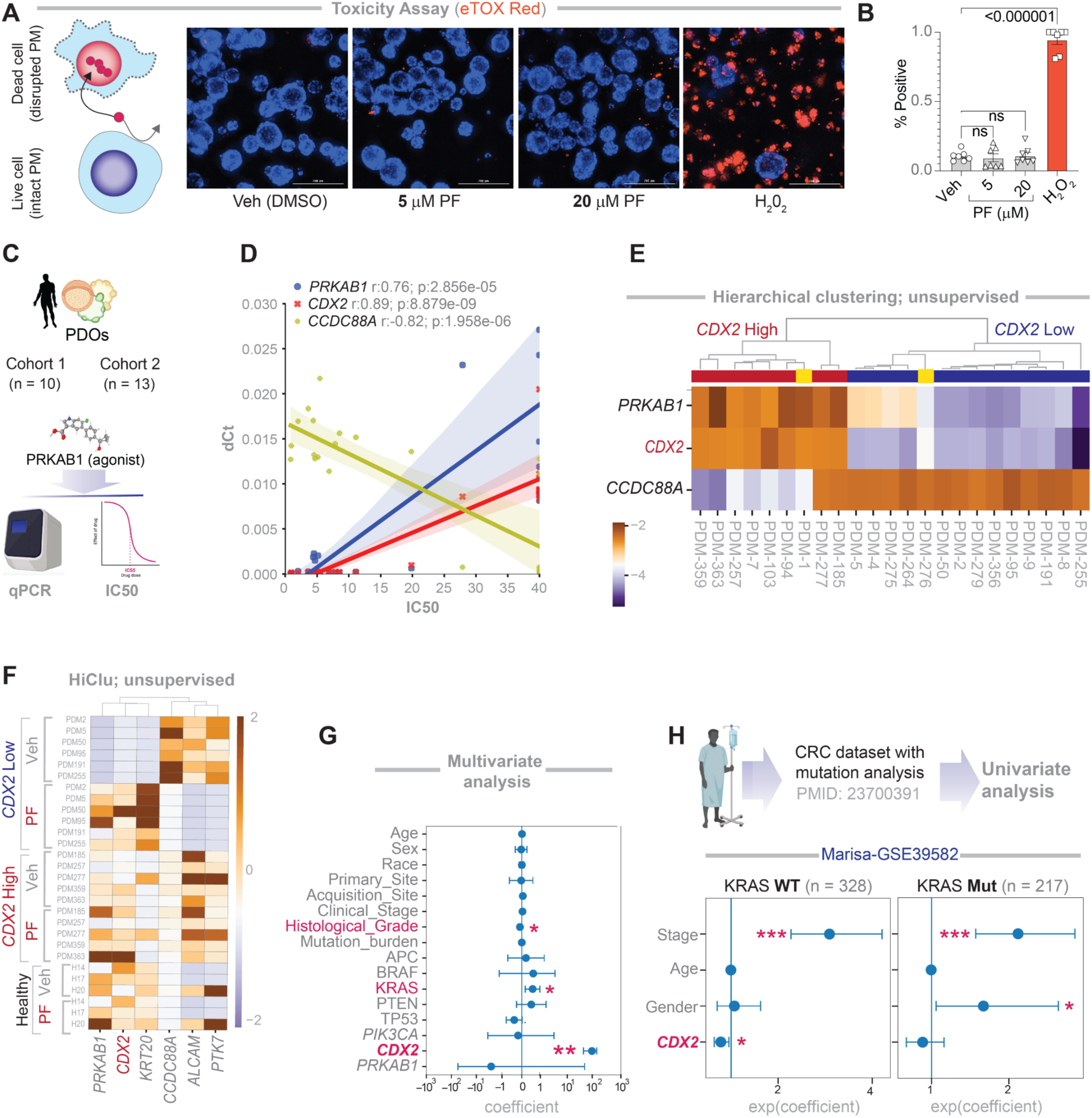
Differentiation and anti-cancer effects of PRKAB1 agonism are restricted to *CDX2*-low PDOs. **A.** Schematic illustrating the principle of eTOX Red staining, which selectively labels dead cells based on increased membrane permeability (left). Representative immunofluorescence images show healthy PDOs treated with vehicle (control), 5 μM PF, 10 μM PF, or H₂O₂ (positive control for cell death) (right). Scale bar = 200 µm. **B.** Quantification of the % of marker-positive (dead) organoids from IF images shown in panel A (right). Statistical comparisons between treatment groups were performed using one-way ANOVA. **C.** Schematic of the experimental workflow showing sequential analyses conducted in two independent patient cohorts. See **Supplemental Information 3** for patient demographics, tumor characteristics and mutation status of key genes. **D.** Correlation plots showing the relationship between IC_50_ values for PF and mRNA expression levels of 3 network-derived genes, as determined by qPCR. **F.** Heatmap showing unsupervised clustering of CRC PDOs based on Z-score-normalized qPCR estimation of the 3 genes in D. **F**. Heatmap comparing expression profiles of stemness and differentiation markers in vehicle-versus PF-treated CRC PDOs. **G**. Multivariate regression analysis of IC_50_ values using a linear model incorporating all measured variables. Bar plot shows the coefficient estimates (center values), 95% confidence intervals (error bars), and p-values for each variable. The p-value for each term tests the null hypothesis that the coefficient is equal to zero (no effect). Red = statistically significant covariates. *p ≤ 0.05; **p ≤ 0.01. See **Supplemental Information 3** for source data. **H**. Univariate analysis of relapse-free survival (RFS) stratified by KRAS mutation status (WT vs. mutant) using dataset GSE39582. *p ≤ 0.05; **p ≤ 0.01; ***p ≤ 0.001.

Using only these three network-derived genes (*CDX2, PRKAB1*, and *CCDC88A*), hierarchical unsupervised clustering accurately classified sensitive versus resistant PDOs (**Figure 3E**). As in prior models, PF promoted differentiation and suppressed stemness markers in CRC PDOs (**Figure 3F**) —effects that were again restricted to CDX2-low tumors, consistent with findings from CRC cell lines (**Figure 2C**) and xenografts (**Figure 2K**).

Multivariate analysis of CRC PDOs (see **Supplemental Information 3** for covariates) identified CDX2 expression as the most significant predictor of IC₅₀ (**Figure 3G**). Two additional variables also emerged: histological grade and *KRAS* mutation status. Specifically, PF was less effective in moderately or well-differentiated tumors that more frequently harbor KRAS mutations (**Figure 3G**). The interplay between CDX2 induction and *KRAS* status suggested a potential ceiling effect—i.e., CDX2 may remain constitutively high in *KRAS*^mut^ tumors, limiting the therapeutic benefit of PF. A univariate analysis in a large CRC cohort with known driver mutations and patient outcomes confirmed this relationship: CDX2 was prognostic in *KRAS*^WT^ tumors (**Figure 3H***-left*), but not in *KRAS*^mut^ tumors (**Figure 3H***-right*).

These findings indicate that despite its modest size, our PDO cohort captures clinically relevant heterogeneity and reveals a key interaction between *CDX2* status and *KRAS* mutation in determining therapeutic response.

### Indicators of therapeutic efficacy and target cell specificity

RNA sequencing across all three model systems—CRC cell lines, xenografts, and PDOs—confirmed that PRKAB1 agonism met the three a priori defined therapeutic success metrics: (i) reinstatement of CDX2, the primary therapeutic goal (bubble plots in **Figure 4A***-top row*; **4B**; violin plots in **Figure S6A-C**), (ii) induction of markers for crypt-top enterocytes and goblet cells, established surrogates of differentiation in the colon crypt^51,85^ (**Figure 4A***-middle row*; **Figure S6D-F**), and (iii) predictable perturbation of the network, specifically induction of genes in clusters #2 and #3 (**Figure 4A***-bottom row*; **Figure S6G-I**). Importantly, these effects were observed exclusively in *CDX2*-low CRC models—cell lines, xenografts, and PDOs—but not in *CDX2*-high PDOs or healthy colon PDOs (**Figure 4A-***columns #5 and #6*; **Figure S6C, S6F, S6I**).

**Figure 4.**
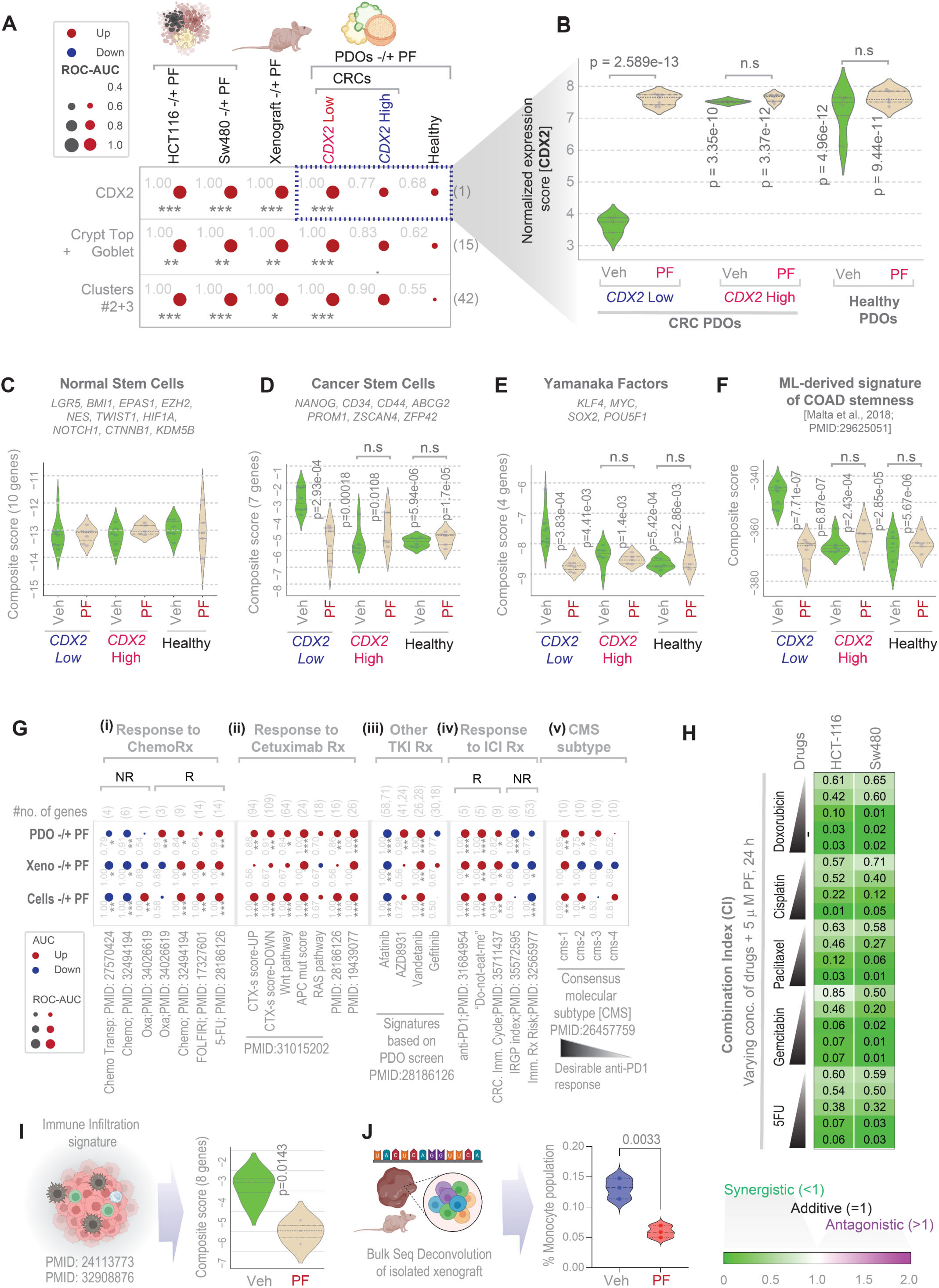
Indicators of therapeutic success, target-cell specificity, and potential for synergy with other treatment modalities. **A**. Bubble plots display ROC-AUC values for the classification of vehicle (Veh) vs. PF-treated samples across multiple models (HCT116, SW480, xenografts, and PDOs). Shown are *CDX2* expression patters (top row), crypt-top and goblet cell markers (middle row), and genes from network clusters #2–3 (bottom row). Circle size reflects the magnitude of the ROC-AUC; color indicates direction of regulation in treated samples (red = upregulated, blue = downregulated). Statistical significance was calculated using Welch’s t-test and annotated using standard codes (‘’p > 0.1; ‘.’p ≤ 0.1; *p ≤ 0.05; **p ≤ 0.01; ***p ≤ 0.001). See **Figure S7** for the same data shown as violin plots with actual p-values. **B**. Violin plots show *CDX2* expression levels in PDOs. Comparisons are made against vehicle (Veh)-treated *CDX2*-low PDOs using Welch’s t-test. **C-F.** Violin plots display the composite score of stemness markers derived from a recent pan-cancer machine learning study^125^. Comparisons were made against vehicle (Veh)-treated *CDX2-*low PDOs using Welch’s t-test. **G.** Bubble plots display ROC-AUC values for gene signatures associated with response (default or indicated with “R”) and non-response (“N.R.”) to multiple therapeutic modalities, across CRC cell lines, xenografts, and PDOs. Circle size reflects ROC-AUC; colors indicate direction of gene regulation. Significance was assessed using Welch’s t-test and annotated as in panel A. **H.** Heatmap shows combination index (CI) values derived from MTT assays in HCT116 and SW480 cell lines treated with a fixed concentration of PF (5 µM, 24 h) and increasing doses of chemotherapeutic agents. CI values were calculated using a previously established formula^126^. **I.** Violin plot shows the composite score of an 8-gene immune infiltration signature^127^ [derived from a large pan-cancer study^128^ (https://bioinformatics.mdanderson.org/estimate/) after training on outcome-annotated datasets] in PF-treated and vehicle (Veh) treated xenografts. p-value was derived using Welch’s t-test. **J.** Violin plot shows the % of monocyte population in PF-treated vs. vehicle (Veh)-treated xenografts, estimated via bulk RNA-seq deconvolution. p-value was derived using Welch’s t-test.

A recent study using a novel machine learning algorithm defined transcriptomic and epigenetic signatures that distinguish normal stem cells from their differentiated progeny, unveiling CSC-specific gene sets indicative of oncogenic dedifferentiation^86^. Applying these signatures, we found that CDX2-reinstatement therapy did not affect normal stem cell–related genes in either healthy or CRC PDOs (**Figure 4C**). In contrast, it selectively downregulated: (i) genes associated with cancer-related stemness (**Figure 4D**), (ii) Yamanaka factors that promote somatic cell reprogramming and are implicated in CSC origins^87^ (**Figure 4E**), and (iii) CRC-specific CSC gene signatures (**Figure 4F**).

These findings demonstrate that *CDX2*-reinstatement acts as a trigger for broader transcriptomic reprogramming that selectively suppresses CSC programs unique to *CDX2*-low tumors, while sparing normal stem cells in healthy colon and *CDX2*-high tumor cells.

### Potential for synergy with other therapeutic modalities

CDX2-low Stage II–III CRCs have previously been shown to derive benefit from adjuvant chemotherapy, whereas CDX2-high tumors do not^52^. We therefore asked whether therapeutic reinstatement of CDX2 in CDX2-low tumors would enhance responsiveness to chemotherapy. Across all models tested, PF treatment consistently altered patient-derived gene expression signatures—inducing those associated with favorable response and suppressing those linked to resistance to various chemotherapeutics (e.g., 5-FU, Oxaliplatin, FOLFIRI) (**Figure 4G-i**). In line with these transcriptional changes, PF (5 µM, 24 h) exhibited synergy with multiple chemotherapeutic agents (see **Figure 4H** legend).

We extended this analysis to other treatment modalities. PF similarly induced gene signatures predictive of response to anti-EGFR therapy (Cetuximab; **Figure 4G-ii**) and several tyrosine kinase inhibitors (TKIs; **Figure 4G-iii**), with a few notable exceptions (Afatinib and Gefitinib; **Figure 4G-iii**). It also enhanced signatures associated with immunotherapy response while suppressing those of non-response (**Figure 4G-iv**). Consistent with this, PF robustly induced the consensus molecular subtype CMS1 across all three model systems (**Figure 4G-v**). This subtype is characterized by a favorable immune landscape and enhanced sensitivity to immune checkpoint inhibitors^88^. In vivo xenograft studies also revealed a treatment-associated reduction in the consensus molecular subtype CMS4, which is linked to poor prognosis and characterized by high stromal content and TGF-β signaling^88,89^. Consistent with this, PF treatment significantly reduced stromal infiltration, as determined by two independent gene expression signatures (see **Figure 4I-J** legend).

These findings suggest that *CDX2*-reinstatement acts as a trigger for broader transcriptomic reprogramming that favors existing treatment modalities, including chemotherapy, targeted therapies, and immunotherapy.

### A molecular signature of therapeutic response and selectivity

To characterize the therapeutic response to CDX2-reinstatement and its impact on the broader transcriptome, we performed integrated differential expression analysis using RNA-seq data from all three models (CRC cell lines, xenografts, and PDOs; see *Methods*). Differentially expressed genes (DEGs) from two independent models (cell lines and xenografts; **Supplemental Information 4-5**) were refined using data from the third (CRC PDOs), yielding a 50-gene signature (**Figure 5A**). These genes were significantly upregulated in *CDX2*-low CRC PDOs compared to healthy and *CDX2*-high CRC PDOs (**Figure 5B**; **Figure S7A-B**). Upon PF treatment, this signature was downregulated to levels comparable to healthy and *CDX2*-high CRC PDOs (**Figure 5B**; see **Supplemental Information 6**), suggesting a shift from a stem-like to a differentiated state.

**Figure 5.**
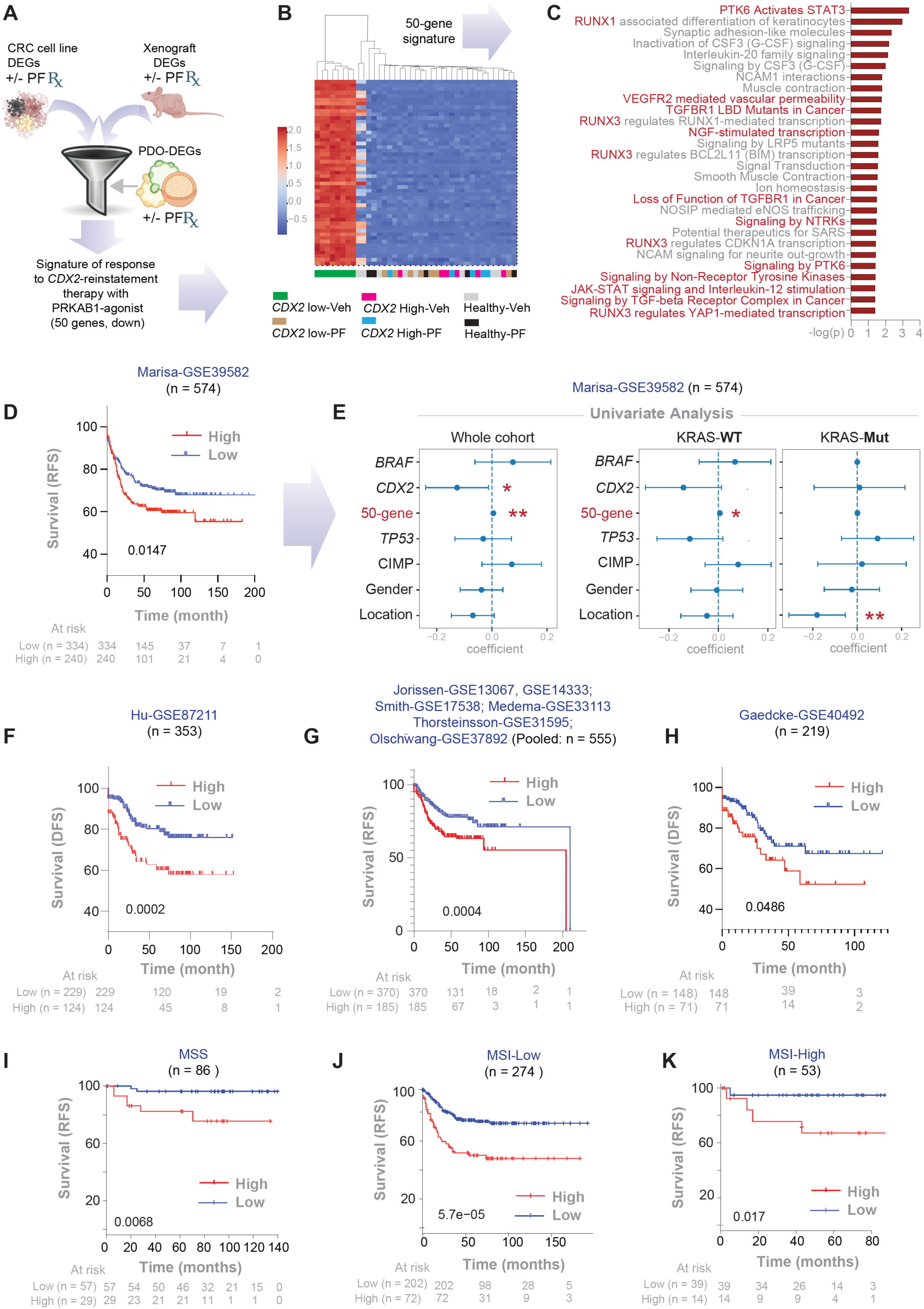
A 50-gene signature for tracking therapeutic response and estimating impact on survival. **A.** Computational workflow used to derive a 50-gene signature of therapeutic response through an integrated differential expression analysis (DEA) approach. Differentially expressed genes (DEGs) from CRC cell lines and xenografts (see **Supplemental Information 4-6**) were further refined based on their ability to classify accurately PDO samples treated with or without PF (see **Supplemental Information 6** for gene list). **B.** Heatmap shows the results of unsupervised clustering of PDOs based on the 50-gene signature. Veh, vehicle. See **Figure S8A-B** for violin plot visualizations. **C.** Reactome pathway analysis of the 50-gene signature, highlighting key biological processes. See **Figure S8C** for enrichment of KEGG pathways. **D**, **F-H**. Kaplan-Meier (KM) plots show recurrence-free (RFS) or disease-free (DFS) survival in various CRC patient cohorts, stratified by high vs. low expression scores of the 50-gene signature based on *StepMiner* algorithm^82^ computed within each cohort. p-values were calculated using the Log-rank test. For comparison, see **Figure S8C** for KM plots on the same cohorts based on *CDX2* expression, computed using the *StepMiner* algorithm^82^ within each cohort. **I-K.** KM plots show recurrence-free (RFS) survival in all *KRAS*^WT^ CRC patients available in KM-plotter database^129^, grouped by their microsatellite stability status. The patient groups are stratified by high vs. low expression scores of the 50-gene signature based on *auto selected best percentile cutoff* algorithm used in KM plotter^129^. p-values were calculated using the Log-rank test. **E**. Univariate Cox regression analysis of RFS in a large CRC cohort. Plots display the coefficient for each variable (center point) with corresponding 95% confidence intervals (error bars) and associated p-values. The analysis includes the 50-gene signature along with other clinical variables. Left: entire cohort; middle: KRAS wild-type CRCs; right: KRAS mutant CRCs. *p ≤ 0.05; **p ≤ 0. 01.

This 50-gene signature included several unique cancer stem cell (CSC)-associated genes—*AGR2*^90^, *ASCL2*^91,92^, *ALDH3A1*^93^, and *TM4SF1* ^94,95^ —not previously captured in machine-learning-derived CSC signatures (**Figure 4C-F**). Pathway enrichment analysis revealed activation of pro-survival programs known to sustain CSCs^96–99^, including PTK6-dependent JAK/STAT signaling, growth factor receptor and non-receptor tyrosine kinases, Hippo/YAP signaling, and TGFβ suppression (**Figure 5C**). Notably, these pathways were selectively downregulated in *CDX2*-low CRCs, but not in healthy or *CDX2*-high CRCs. Similar pathway enrichment analyses of the upregulated genes showed genes associated with epithelial structure maintenance (**Figure S7C**).

### Impact of CDX2 reinstatement therapy on survival

To evaluate clinical relevance, we tested whether the 50-gene signature could stratify relapse-free/disease-free survival (RFS/DFS) in multiple CRC patient cohorts. Across all datasets, patients with low 50-gene scores—resembling PF-treated or *CDX2*-high tumors—had significantly better outcomes than those with high scores (**Figure 5D, F-K**). The associated odds ratio was ∼2.0 (**Supplemental Information 7-***tab #1*), indicating that elevated expression of the signature doubles the risk of recurrence or death—risk that may be mitigated by CDX2-reinstatement therapy.

Univariate analysis showed that both *CDX2* expression and the 50-gene signature were significant predictors of RFS (**Figure 5E***-left*), but the signature performed better, especially in *KRAS*^WT^ tumors (**Figure 5E***-middle*). Across all patient cohorts, the 50-gene score consistently outperformed *CDX2* as a prognostic indicator (**Figure S7D-G**; **Supplemental Information 7**). As seen previously for *CDX2* (**Figure 3E**), the 50-gene signature did not correlate with outcome in *KRAS*^mut^ tumors (**Figure 5E***-right*), suggesting that *KRAS* mutation may bypass the CDX2 axis. In fact, within the *KRAS*^WT^ group, patients with low 50-gene scores resembling PF-treated tumors had significantly better outcomes than those with high scores regardless of microsatellite stability status (**Figure 5J-L**).

Together, these findings identify a robust, target-derived 50-gene signature that not only captures the molecular response to *CDX2*-reinstatement but also stratifies survival risk. Therapeutic suppression of this signature emerges as a measurable and meaningful objective, with potential to improve patient outcomes.

### State-specific cell fates dictate selectivity of CDX2-restorative therapy

Hallmark pathway analysis revealed that *CDX2*-restoration via PF treatment consistently activated a transcriptional program across all three models centered on the restoration of epithelial integrity, including polarity and cell junction assembly (**Figure 6A**). Integration of downregulated genes across models identified a core set of commonly suppressed transcripts (**Figure 6B**), and transcription factor (TF) enrichment analysis implicated key stemness-associated TFs, e.g., *TEAD4*, *SP1*, and *KLF1/4* as top regulators of these genes (**Figure 6C**).

**Figure 6.**
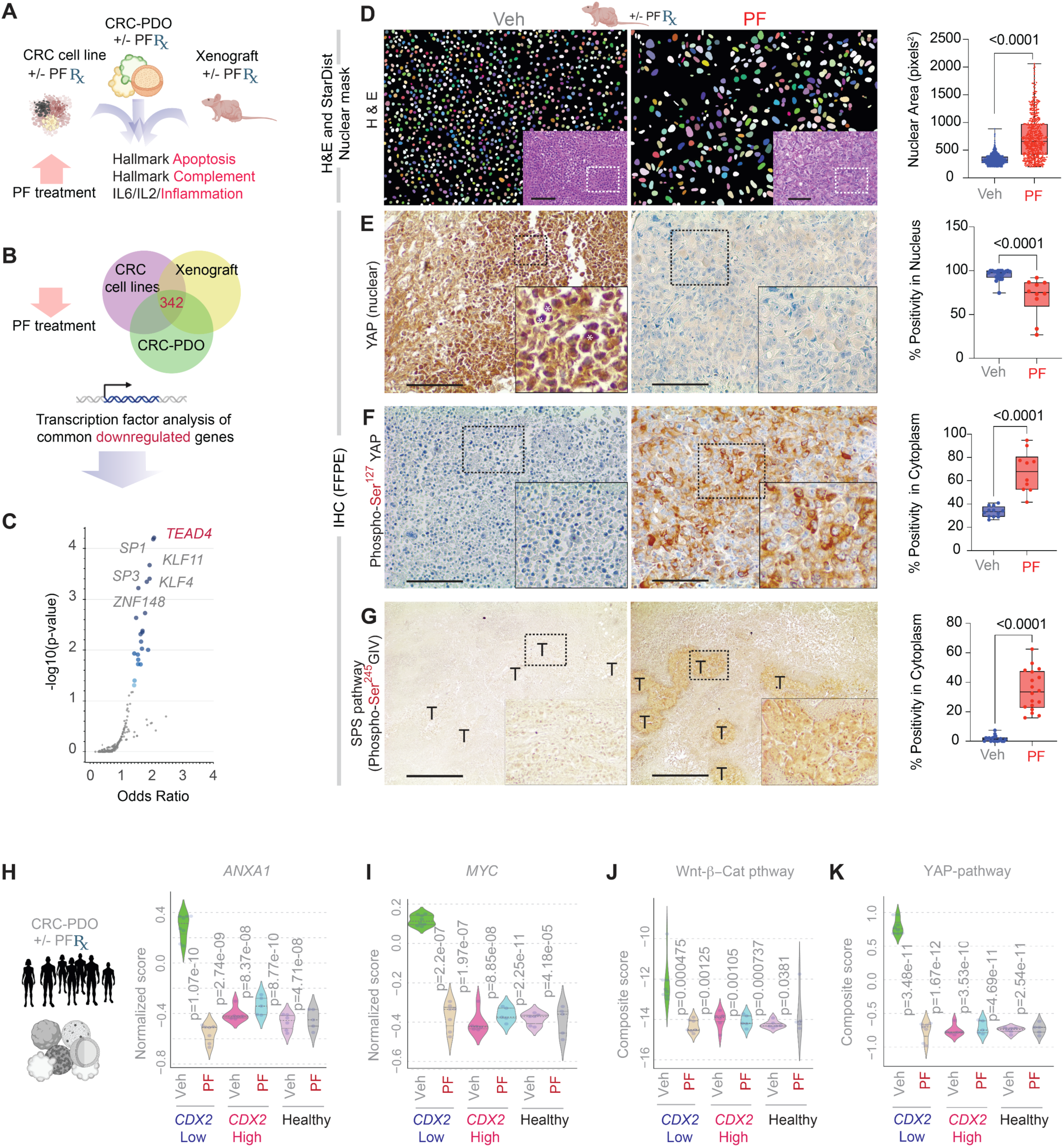
CDX2-reinstatement reprograms CRC stem cells by restoring epithelial polarity. **A.** Schematic summarizes top hallmark pathways identified using integrated differential expression analysis (DEA) approach that are induced upon PF treatment in three model systems. **B-C.** Venn diagram (B) depicts downregulated genes common in all the three models used in this work. Transcription factor (TF) analysis (C) identified top TFs associated with common downregulated genes upon PF treatment in all the models. **D.** Representative H&E-stained sections (left, with inset) of FFPE xenograft tumors from vehicle (Veh) and PF-treated groups, with corresponding quantification of nuclear area (right) using StarDist-based segmentation in ImageJ. **E-G**. Immunohistochemistry (IHC) images (left) and quantification (right) of xenografts showing expression of: nuclear YAP (E; white asterisk, inset); phospho-Ser^127^ YAP (F); and phospho-Ser^245^ GIV (a biomarker of the SPS pathway^78–80^; see also **Figure S8A**). Scale bar = 100 µm. **H-K**. Violin plots showing normalized expression scores in vehicle (Veh)- and PF-treated PDOs for transcriptional markers of oncogenic stemness: Anexin (H), MYC (I), composite Wnt/β-catenin (J) and YAP (K) target genes (listed in Figure 7-*left*). p-values were calculated using Welch’s t-test, comparing each sample to vehicle-treated *CDX2*-low PDOs.

Histological analysis of xenograft tumors reinforced these transcriptional findings. PF-treated tumors displayed a significantly increased number of asymmetrically stretched nuclei, consistent with chromatin reorganization during epithelial reprogramming (**Figure 6D**). These morphological changes were accompanied by pronounced alterations in Hippo signaling: PF reduced nuclear localization of YAP (**Figure 6E**) while increasing its inactive, phosphorylated (p)-Ser^127^ form (**Figure 6F**). This is consistent with pSer^127^-mediated cytosolic retention and phosphodegron-dependent degradation of YAP oncogenic activity^100^, explaining its near-total loss in treated tumors (**Figure 6E**).

PRKAB1 is also known to activate the stress polarity signaling (SPS) pathway^78–80^; it is a sentinel system that integrates metabolic stress with epithelial polarity, coordinating cell–cell junction restoration under bioenergetic challenge (see **Figure S8A**). In normal epithelial cells, SPS is inducible and supports junctional reassembly, anchorage-dependent growth as monolayer, and epithelial barrier integrity^30,79^. In contrast, transformed cells lose this pathway^79^, enabling anchorage-independent growth and junctional disassembly (**Figure S8B-C**). Notably, (re)activation of SPS permits anchorage-independent growth of normal cells but not transformed cells^80^—highlighting its potential to discriminate between cell states and a plausible reason for PF’s ability to selectively target CSCs, but spare healthy stem cells. We found that PF induced the SPS pathway, as evidenced by upregulation of a canonical biomarker pSer^245^GIV^30,79,80^ (**Figure 6G**).

Next, we hypothesized that SPS reactivation preserves the function of normal stem cells while selectively dismantling the survival programs of CSCs, which rely on SPS pathway silencing. To test this, we analyzed expression scores for core transcriptional targets of oncogenic stemness in PF-treated healthy and CRC PDOs. PF significantly reduced the expression of *Annexin A1* (**Figure 6H**), *MYC* (**Figure 6I**), and transcriptional programs of Wnt/β-catenin (**Figure 6J**) and YAP (**Figure 6K**)—all of which were elevated in *CDX2*-low CSC-rich PDOs but not in healthy- or *CDX2*-high CSC-poor PDOs.

Together, these findings demonstrate that CDX2-reinstatement reprograms CSCs by restoring epithelial polarity through SPS pathway reactivation. This reactivation triggers the collapse of two key junction-informed transcriptional programs—YAP and Wnt/β-catenin—on which CDX2-low CSCs are critically dependent (see **Figure 7A-B** legend). This mechanism underpins the selectivity and efficacy of PF, driving differentiation in all cells but apoptotic fate specifically in CSCs (**Figure 7C**).

**Figure 7.**
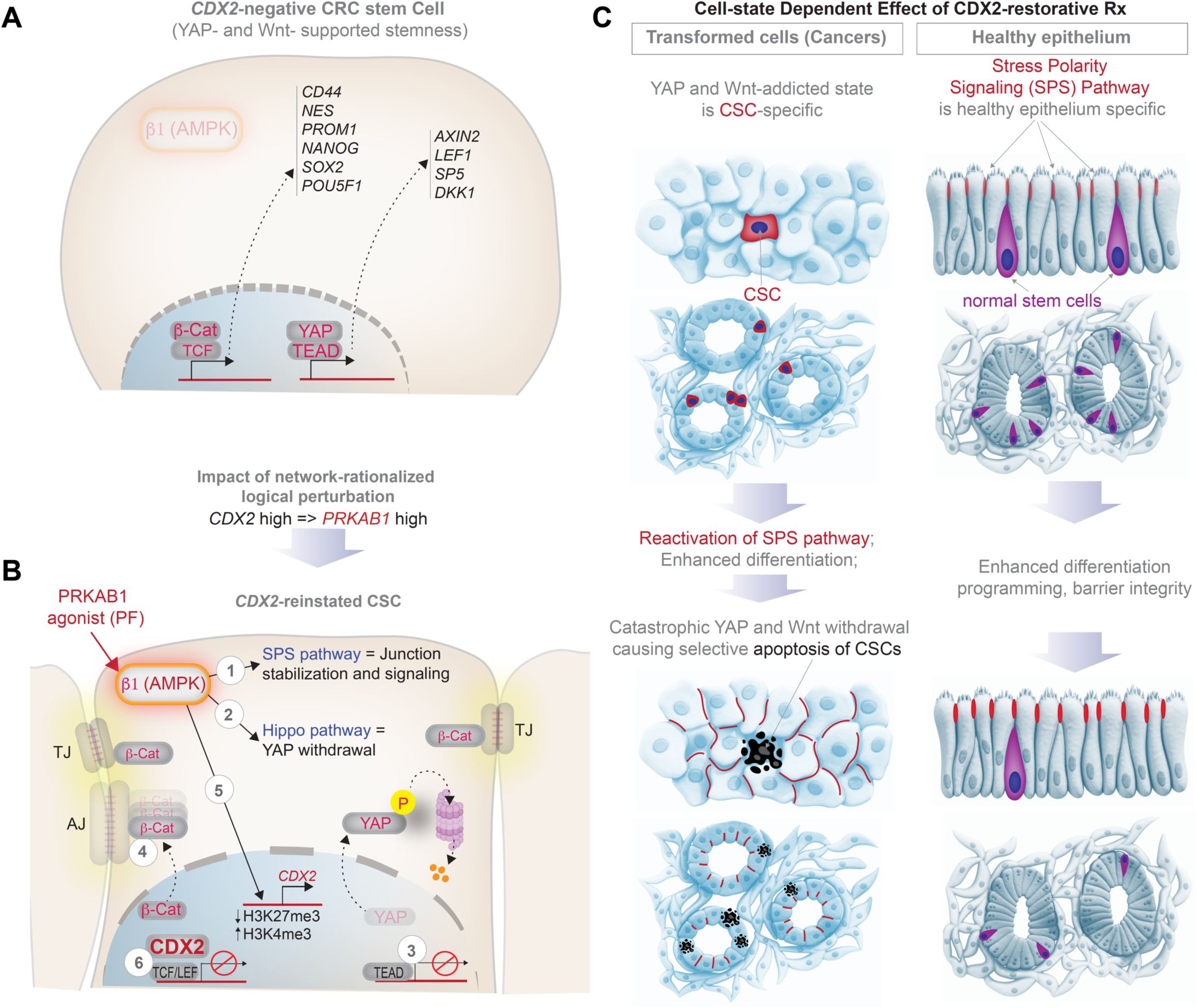
Summary and working model of how CDX2-reinstament selectively targets CRC stem cells. **A.** CRC stem cells, but not normal stem cells, are characterized by enhanced transcription of Wnt/β-catenin and YAP target genes that are established markers of cancer stem cells. Both pathways are known to collaboratively enhance stemness in the colon^130,131^. **B.** PF treatment activates β1-specific AMPK, reactivates a tumor suppressor pathway (SPS pathway; 1) which attempts to restore tight-(TJ) and adherens (AJ) junctions in the setting of bioenergetic stress. Bioenergetic stress (AMPK activation) and junction restoration reshapes two junction-informed stemness programs, leading to their catastrophic withdrawal: *(i)* the Hippo-YAP pathway is *activated* (2), resulting in YAP phosphorylation followed by proteasomal degradation, and consequently, its nuclear exclusion (3) and withdrawal of the YAP/TEAD transcriptional program; *(ii)* junctional sequestration of β-Catenin and consequently, its nuclear exclusion (4) and termination of the β-Catenin•TCF/LEF transcriptional program. *CDX2*-reinstatement, and differentiation programs that accompanies the same, is likely via AMPK-dependent epigenetic events^123,124^ (5). Once translated into protein, CDX2 is known to complete and displace β-catenin from TCF/LEF complexes^132^ (6), which further accentuates the impact of the SPS pathway on the β-Catenin•TCF/LEF transcriptional program. **C.** Schematic illustrates the cell state-specific impact of CDX2-reinstatement therapy and the impact of restoration of the stress polarity signaling (SPS) pathway. Pre-(top) and post-(bottom) treatment states of cancers (left) and healthy (right) epithelium are shown. Although activation of the SPS pathway enhances differentiation and junction formation in both cell states, the apoptotic fate is selectively seen in *CDX2*-low CSCs.

## Discussion

The central discovery of this study is a network-guided differentiation therapy that reinstates CDX2 expression to selectively eliminate CSCs in CRCs. Much like immunotherapy reawakens immune surveillance, CDX2-restorative therapy reactivates a physiologic lineage commitment. We identify *PRKAB1* as a key upstream node and show that PF-06409577, a clinically safe first-in-class agonist, triggers *CDX2*-restoration, cell-type specific differentiation and anti-tumor responses. The therapeutic effect meets rigorous, pre-defined benchmarks for efficacy: *CDX2* induction, a robust 50-gene companion biomarker signature, suppression of stemness and survival pathways, and sparing of normal stem cells. Additional analyses of human CRCs using a panel of treatment-perturbed genes promises a ∼50% reduction in relapse and mortality risk in *CDX2*-low CRCs.

### A Scalable Framework for Differentiation Therapy

Our use of the BoNE enabled the development of *CANDiT*, a rational, systems-level framework for inducing differentiation in solid tumors. By using *CDX2*—a master regulator of intestinal lineage—as a seed node, BoNE traced upstream regulatory hubs capable of reinstating epithelial identity while suppressing CSC-associated programs in colorectal cancers. This approach is generalizable: master regulators of differentiation in other tissues can similarly be used to drive lineage restoration, examples of which are emerging^101^.

Although *CDX2* is insufficient to fully activate the lineage program^75^, our findings suggest that PRKAB1 agonism by PF enables broader commitment by sustaining *CDX2* induction and engaging additional, underexplored and incompletely mapped pathways essential for terminal differentiation. *CANDiT* distinguishes itself from conventional screening methods by revealing hidden transcriptomic vulnerabilities in dedifferentiated CSCs and pinpointing high-value targets to exploit them. In the PRKAB1-activated state, *CDX2* plays a central—but not solitary—role in driving differentiation and CSC-specific death. Like *CDX2*, *PRKAB1* is likely to be necessary, but not sufficient to impose such broader commitment to differentiation. Our findings exemplify how *CANDiT* offers a scalable, mechanistically grounded blueprint for developing lineage-based therapies across solid tumors, guided by tissue-specific master regulators.

### Mechanism of Selectivity

We show that CDX2-restorative therapy exploits a selective vulnerability in CSCs by reactivating the SPS pathway, a circuit integrating bioenergetic stress with epithelial polarity in 3D growth contexts. While this pathway is silenced in cancer, it remains functional in healthy tissues^78,79^. PRKAB1 agonism restores epithelial polarity through SPS activation, disrupting anchorage-independent growth and silencing junction-informed oncogenic programs (e.g., YAP, Wnt/β-catenin; **Figure 7A-B**), leading to a collapse of stemness signaling in CSCs. In contrast, in normal epithelial and stem cells, SPS activation supports barrier function and preserves homeostasis. This cell-state–specific response explains the therapeutic selectivity of PF, enabling CSC-targeted killing while sparing healthy stem cells (see **Figure 7C**).

It is noteworthy that the Cancer Dependency Map (DepMap; https://depmap.org/portal/home/) includes data for HCT116 and SW480, but not for key lines used here such as DLD1 and Caco2. Among the cell lines represented, PF IC₅₀ values do not correlate with CDX2 transcript abundance. Our mechanistic data— highlighting the role of 3D context and anchorage-independent signaling in PF response— offer a plausible explanation, aligning with DepMap’s own push to include 3D IC₅₀ datasets^102^.

### Translational Potential and Clinical Prioritization

We advocate translating these findings into clinical trials with a focus on CDX2-low CRCs, especially those with a high 50-gene response signature. CDX2-low status, as assessed by IHC, is prognostic in stage III/IV disease and is frequently associated with *BRAF* mutations^103^. Conversely, *KRAS*^mut^ tumors tend to retain CDX2, reflecting the role of *KRAS^mut^* in promoting hyperplasia^104^ rather than stemness^105^. This intriguing distinction could be attributed to cell-type specific roles of *KRAS*/*BRAF*^mut^ reported recently^106^.

However, KRAS status alone should not preclude PF therapy: both HCT116 and SW480 cells (CDX2-low and *KRAS*^mut^) were PF-sensitive. Notably, population studies confirm that a subset (∼9%) of *KRAS*^mut^ metastatic CRCs with low CDX2 have poor prognosis^103^ and could benefit from differentiation therapy. Thus, CDX2-low status and high therapeutic response signature, not KRAS or BRAF status, should drive clinical candidacy. Because loss of *CDX2* is an adverse prognostic factor in stage IV and stage III/*BRAF*^mut^ CRCs^74,107^, these subgroups can be prioritized for *CDX2*-reinstatement therapy with PRKAB1 agonist as single agent or together with other modalities.

Although *CDX2* is prognostic in stage II(MSS)^52,65,69^ and stage III CRCs^52,108^, caution is warranted when combining PF with conventional chemotherapy. CDX2 expression is associated with MDR1 upregulation, which could contribute to chemoresistance^109,110^. Until further data are available, PF should be prioritized as monotherapy or in combination with non-cytotoxic agents in patients with relapsed or metastatic *CDX2*-low CRC. Finally, early-onset CRCs is a subgroup that show worse outcome (poorer DFS) than its late-onset counterparts^111^. Because the *CDX2*-*PRKAB1* relationship is conserved in this subgroup (**Figure S2A**), we propose including this group in future PF-focused trials.

### Implications for Pre-Neoplastic Disease

It has been recently shown that CDX2 loss plays a permissive role in CRC initiation, particularly along the serrated pathway^112–114^ and in inflammation-associated CRCs^115–117^. Mouse models have shown that *CDX2*-deficient metaplastic cells can induce tumorigenesis non–cell-autonomously^118,119^. Our findings provide an impetus for future studies to evaluate whether *PRKAB1* agonists can serve as prophylactic agents in high-risk populations with pre-neoplastic lesions or chronic inflammation.

## Conclusions

We present *CANDiT*, a machine learning–guided, network-based differentiation strategy that fills a critical gap in cancer therapy. *CANDiT* differs fundamentally from traditional pipelines in three key ways: 1) Network-transcriptomics and ML-guided target discovery; 2) Validation studies on network-vetted cell line and xenograft models; 3) Validation in PDOs utilizing biomarker-driven stratification and adjusting for confounding variables as done in clinical trial analysis. This strategy enables selective CSC targeting and spares normal stem cells. Despite strong evidence for CDX2 as a prognostic biomarker^52,65,66,68,69,71,83,120,121^, and a compelling pharmacoeconomic justification (i.e., cost $ 50 000 per QALY) for a CDX2-stratified decision to offer chemotherapy^122^, its clinical utility has not been adopted into guidelines due to lack of prospective validation. Our demonstration that CDX2 serves as both a predictive biomarker and therapeutic effector for PF—a drug already deemed safe in humans (NCT02286882)— provides compelling justification for CDX2-guided clinical trials. More broadly, *CANDiT* opens the door to rational, scalable differentiation therapy across solid tumors.

## Limitations of the study

The exact mechanism(s) linking PRKAB1, SPS activation, *CDX2* restoration and crypt differentiation was not fully delineated. Future studies will examine whether SPS (re)activation reverses aberrant *CDX2* promoter methylation^123,124^. The PDO cohort was limited in size but reflective of real-world diversity, capturing the interplay between tumor differentiation, CDX2 expression, and *KRAS*^mut^ status; larger cohorts and randomized clinical trials with RNA-seq endpoints is warranted. Epigenetic profiling and cell fate trajectory analysis will be essential to uncover the basis for the divergent cell fates—differentiation versus apoptosis— triggered by PF in normal vs CSCs.

## Supporting information

SOM

## Acknowledgements

We are grateful to the staffs of the UC San Diego HUMANOID™ Center of Research Excellence, for support in the logistical support in conducting the studies on human PDOs. This work was supported by the National Institutes for Health (NIH) grants UG3TR002968, UH3 TR002968, R01-CA238042, R01-CA100768 and R01-CA160911 (to P.G) and VA Merit awards 1 I01 BX003856-01A1 and 1 I01 BX004494-01 (M.B). S.S was supported in part by a American Association of Immunologists (AAI) Intersect Fellowship Program for Computational Scientists and Immunologists. SB was supported by SERB International Research Experience Fellowship (SIR/2022/001374) by Govt. of India. Other sources of support include: R01-AI155696, R01-AI141630 and UG3TR003355 (P.G), a Torrey Coast Foundation Award (P.G, M.B, J.Y), Curebound Foundation award 23DG04, and by three Padres Pedal the Cause awards #PTC2021, #PTC2022 and #PTC2024. P.G were also supported by the Leona M. and Harry B. Helmsley Charitable Trust. Funders had no role in study design or conclusions. Authors acknowledge the instrumentation resources at the US San Diego Agilent Center of Excellence in Cellular Intelligence and Debashis Sahoo (UC San Diego) for providing access to computational resources.

## Author Contributions

SS and PG conceptualized the study; PG designed and analysed all parts of the study; JA, KP, AO, EV and CT acquired, expanded, biobanked and conducted all experiments with patient-derived organoids and CRC cell lines; VC conducted immunocytochemistry (IHC) and immunofluorescence (IF) and Hematoxylin and Eosin (H&E) staining studies; CRE conducted qPCR studies and assisted SS, VC and PG on key parts of the study; SB, EM, KP, SS and VC conducted IHC, IF and H&E quantifications. ST built and validated the network model; SS and PG conducted all other computational, statistical and bioinformatic analyses; SA and MB conducted the xenotransplantation studies; JRS and JY synthesized the pharmacophore; MB, JY and PG secured funding; SS and PG prepared all display items; PG wrote the first draft of the manuscript; all authors edited and approved the final version.

## Star Methods

### Key Resource Table

| REAGENT or RESOURCE | SOURCE | IDENTIFIER |
| --- | --- | --- |
| **Biological Samples and Cell Lines** |  |  |
| HCT-116 | ATCC | CCL-247 |
| Sw480 | ATCC | CCL-228 |
| DLD1 | ATCC | CCL-221 |
| Caco2 | ATCC | HTB-37 |
| *Athymic nude male mice* | Jackson Laboratory | 002019 |
| PDM-255 | ATCC | HCM-CSHL-0245-C18-A |
| PDM-2 | ATCC | HCM-CSHL-0057-C18 |
| PDM-4 | ATCC | HCM-CSHL-0060-C18 |
| PDM-50 | ATCC | HCM-SANG-0273-C18 |
| PDM-255 | ATCC | HCM-CSHL-0245-C18-A |
| PDM-275 | ATCC | HCM-CSHL-0245-C18-B |
| PDM-1 | ATCC | HCM-CSHL-0056-C18 |
| PDM-279 | ATCC | HCM-CSHL-0384-D37 |
| PDM-185 | ATCC | HCM-CSHL-0238-C18 |
| PDM-191 | ATCC | HCM-CSHL-0246-C19 |
| PDM-9 | ATCC | HCM-CSHL-0065-C20 |
| PDM-264 | ATCC | HCM-CSHL-0382-C19 |
| PDM-94 | ATCC | HCM-CSHL-0141-C18 |
| PDM-356 | ATCC | HCM-CSHL-0257-C18 |
| PDM-7 | ATCC | HCM-CSHL-0063-C18 |
| PDM-257 | ATCC | HCM-CSHL-0248-C19 |
| PDM-363 | ATCC | HCM-CSHL-0426-C18 |
| PDM-276 | ATCC | HCM-CSHL-0461-D12 |
| PDM-277 | ATCC | HCM-CSHL-0247-C18 |
| PDM-359 | ATCC | HCM-CSHL-0317-C18 |
| PDM-5 | ATCC | HCM-CSHL-0061-C18 |
| PDM-103 | ATCC | HCM-CSHL-0164-C20 |
| PDM-8 | ATCC | HCM-CSHL-0064-C18 |
| PDM-95 | ATCC | HCM-CSHL-0142-C18 |
| Human colon-derived healthy organoids | UC San Diego HUMANOID™ Center | n/a |
| **Chemicals and Reagents** |  |  |
| eTox red | Agilent | Cat#8711009 |
| Guava Cell Cycle Reagent | Millipore Sigma | Cat#SKU4500-0220 |
| Dead Cell Apoptosis Kits with Annexin V | Thermo Fisher Scientific | Cat# V13242 |
| TRIzol™ Reagent | Life Technologies | Cat#15596018 |
| Quick-RNA MiniPrep Kit | Zymo Research | Cat#R2052 |
| qScript™ cDNA SuperMix | QuantaBio | Cat#101414 |
| PowerUp™ SYBR™ Green Master Mix | Applied Biosciences | Cat#A25741 |
| Direct-zol RNA Miniprep Kit | Zymo Research | Cat#R1051 |
| Zinc Formalin Fixative | Sigma-Aldrich | Cat#Z2902 |
| DMEM/High glucose with L-glutamine; without sodium pyruvate | Cytiva | Cat#SH30022.FS |
| Fetal Bovine Serum | Sigma-Aldrich | Cat#F8067-500ML |
| Penicillin Streptomycin 100X Solution | Cytiva | Cat#SV30010 |
| Trypsin-EDTA (0.25%), phenol red | Fisher Scientific | Cat#25-200-114 |
| Dulbecco's Phosphate Buffered Saline (DPBS), 1X, without calcium, magnesium | Cytiva | Cat#16750-076 |
| Matrigel | Corning | Cat#354234 |
| Prochlorperazine dimaleate salt | Selleck Chemicals | Cat#S4631 |
| Mayer's hematoxylin | BioGenex | Cat#HK1005K-GP |
| Two-component DAB brown | BioGenex | Cat#HK542XAKE-GPE |
| **Collagenase I** | GIBCO | Cat#17100-017 |
| **DNase I** | Zymo Research | Cat#E1011-A |
| **Human TruStain FcX™** | BioLegend | Cat#422302 |
| **Poly-L-lysine coated slides** | Fisher Scientific | Cat#63410-02 |
| **3% hydrogen peroxide** | Fisher Scientific | Cat#S25359 |
| **Fluid Thioglycollate Medium** | BD Difco | Cat#225650 |
| **2% paraformaldehyde** | Electron Microscopy Services | Cat#15700 |
| **0.2% gluteraldehyde** | Electron Microscopy Services | Cat#16200 |
| **0.1% Probumin® BSA** | Millipore Sigma | Cat#820452 |
| **2% gelatin** | Sigma-Aldrich | Cat# ES-006-B |
| **Protease inhibitor cocktail** | Roche Diagnostics | Cat#5056489001 |
| **Glutathione Sepharose beads** | Sigma-Aldrich | Cat#17-0756-04 |
| **RPMI 1640 medium** | Thermo Scientific | Cat#11875093 |
| **Gibco TrypLE express enzyme (1X)** | Thermo Fisher Scientific | Cat#12563011 |
| **Intestigro™** | HUMANOID | Cat#HUM2019-125 |
| **Dimethyl sulfoxide (DMSO)** | Sigma-Aldrich | Cat#D2650-100ML |
| **Thiazolyl Blue Tetrazolium Bromide** | Sigma-Aldrich | Cat#M5655-1G |
| **Sodium Dodecyl Sulfate** | Fisher Scientific | Cat#BP166-500 |
| **Dead Cell Apoptosis Kit with Annexin V Alexa Flour 488 & Propidium Iodide (PI)** | Thermo Fischer Scientific | Cat#V13245 |
| **Chemicals and Reagents (For biosynthesis of AMPKβ1 agonist)** |  |  |
| **1,4-Dibromobenzene** | Fisher Scientific | Cat#AC112732500 |
| **Tetrahydrofuran (THF, anhydrous)** | Fisher Scientific | Cat#T425-1 |
| **n-Butyllithium (n-BuLi, 2.5 M in hexane)** | Fisher Scientific | Cat#AC213351000 |
| **Cyclobutanone** | Fisher Scientific | Cat#AC110930250 |
| **Ammonium chloride (NH<sub>4</sub>Cl, saturated aqueous)** | Fisher Scientific | Cat#A2037300G |
| **Ethyl acetate (EtOAc)** | Fisher Scientific | Cat#AA31344M4 |
| **Brine (saturated NaCl solution)** | Fisher Scientific | Cat#S271-10 |
| **Magnesium sulfate (MgSO<sub>4</sub>)** | Fisher Scientific | Cat#M65-3 |
| **Sodium tert-butoxide** | Fisher Scientific | Cat#S0450100G |
| **Potassium acetate** | Fisher Scientific | Cat#P171-500 |
| **XPhos Pd G2 (palladium catalyst)** | Sigma-Aldrich | Cat#741825-1G |
| **XPhos (ligand)** | Sigma-Aldrich | Cat#638064-1G |
| **Tetrahydroxydiboron** | Sigma-Aldrich | Cat#754242-25G |
| **Argon gas** | Matheson Tri-Gas | SKU: AR 200 |
| **5-Bromo-6-chloro-1H-indole** | Combi-Blocks | Cat# QB-9262 |
| **4-Dimethylaminopyridine (DMAP)** | Fisher Scientific | Cat#D1450100G |
| **Pyridine** | Fisher Scientific | Cat#P368-500 |
| **Trichloroacetyl chloride** | Fisher Scientific | Cat#AAB2167518 |
| **Sodium methoxide (25% in methanol)** | Fisher Scientific | Cat#AA46585AP |
| **2-Methyltetrahydrofuran (2-MeTHF)** | Fisher Scientific | Cat#AC396621000 |
| **Potassium carbonate (K<sub>2</sub>CO<sub>3</sub>, aqueous)** | Fisher Scientific | Cat#P208-500 |
| **Sodium hydroxide (NaOH, 3 M aqueous)** | Fisher Scientific | Cat#S318-1 |
| **Hydrochloric acid (HCl, 1 M aqueous)** | Fisher Scientific | Cat#A144-212 |
| **Acetonitrile** | Fisher Scientific | Cat#A998-4 |
| **Dichloromethane (DCM)** | Fisher Scientific | Cat#D37-20 |
| **Trifluoroacetic acid (TFA, 0.1% in water for HPLC)** | Fisher Scientific | Cat#AA44630AP |
| **Antibodies** |  |  |
| **Anti-CDX2 antibody** | Abcam | Cat#76541 |
| **AMPK β1 Polyclonal antibody** | Proteintech | Cat#10308-1-AP |
| **Anti-pSer<sup>245</sup>-GIV antibody** | 21st Century Biochemicals (Marlboro, MA) | Customized |
| **Anti p-YAP antibody** | Cell Signaling Technology | Cat#13008S |
| **YAP antibody** | Santa Cruz | Cat#101199 |
| **Anti-beta catenin** | Santa Cruz | Cat#7963 |
| **Goat anti Rabbit IgG (H+L) Secondary Antibody, Alexa Fluor 647** | Invitrogen | Cat#A-21245 |
| **F(ab')<sub>2</sub>-Goat anti-Rabbit IgG (H+L) Cross-Absorbed Secondary Antibody, Alexa Fluor 594** | Invitrogen | Cat#A-11072 |
| **Software Programs and Algorithms** |  |  |
| **ImageJ** | National Institute of Health | https://imagej.net/Welcome |
| **Prism 9** | GraphPad | https://www.graphpad.com/scientific-software/prism/ |
| **Molsoft** | Molsoft, LLC | https://www.molsoft.com/index.html |
| **LAS-X** | Leica | www.leica-microsystems.com/products/microscope-software/p/leica-las-x-ls |
| **Illustrator** | Adobe | https://www.adobe.com/products/illustrator.html |
| **ImageStudio Lite** | LI-COR | https://www.licor.com/bio/image-studio-lite/ |
| **ClueGO** | Cytoscape | https://academic.oup.com/bioinformatics/article/25/8/1091/324247 |
| **NetworkX** | Python | https://networkx.org |
| **NumPy** | Python | https://numpy.org/ |
| **Pandas** | Python | https://pandas.pydata.org/ |
| **Seaborn** | Python | https://seaborn.pydata.org/ |
| **Gephi** | Gephi | https://gephi.org |
| **Photoshop** | Adobe | https://www.adobe.com/products/photoshop.html |
| **FlowJo** | Flow Jo V10, BD BioSciences | RRID:SCR_008520 |
| **Deposited data** |  |  |
| GSE237624 | Induction of CDX2 as a differentiation therapy in colorectal cancers [cell lines] | This work |
| GSE237625 | Induction of CDX2 as a differentiation therapy in colorectal cancers [xenograft] | This work |
| GSE237623 | Induction of CDX2 as a differentiation therapy in colorectal cancers [CRC organoids] | This work |
| **Other** |  |  |
| **6-well Tissue Culture Plate** | Genesee Scientific | Cat# 25-105 |
| **12-well Tissue Culture Plate** | CytoOne | Cat# CC7682-7512 |
| **Transwell Inserts (6.5 mm, 0.4 μm pore size)** | Corning | Cat# 3470 |
| **Cell Scraper** | Millipore Sigma | Cat# C5981-100EA |
| **Millicell EZ Slide 8-Well Chamber** | Millipore Sigma | Cat# PEZGS0816 |
| **Trypan Blue Stain** | Invitrogen | Cat# T10282 |
| **70 μm Cell Strainer** | Thermo Fisher Scientific | Cat# 22-363-548 |
| **Noyes Spring Scissors - Angled** | Fine Science Tools | Cat# 15013-12 |
| **gentleMACS™ C Tubes** | Miltenyi Biotec | Cat# 130-093-237 |
| **DAPI** | Invitrogen | Cat# D1306<br>RRID: AB_2629482 |
| **Phalloidin, Alexa Fluor 594** | Invitrogen | Cat# A12381<br>RRID: AB_2315633 |
| **Propidium Iodide** | Invitrogen | Cat# V-13245 B |
| **Countess II Automated Cell Counter** | Thermo Fisher Scientific | AMQAX1000 |
| **Epithelial Volt-Ohm (TEER) Meter** | Millipore | MERS00002 |
| **Automated TEER measurement system [REMS AutoSampler]** | World Precision Instruments (WPI) | N/A |
| **MESO QuickPlex SQ 120** | MSD | N/A |
| **Leica TCS SPE Confocal** | Leica Microsystems | TCS SPE |
| **Power Pressure Cooker XL** | Tristar Products | N/A |
| **Canon Rebel XS DSLR** | Canon | N/A |
| **MiniAmp Plus Thermal Cycler** | Applied Biosystems | Cat# A37835 |
| **QuantStudio5** | Applied Biosystems | Cat# A28140<br>RRID:SCR_020240 |
| **Light Microscope (brightfield images)** | Carl Zeiss LLC | Axio Observer, Inverted; 491917-0001-000 |
| **Fisherbrand™ 150 Handheld Homogenizer** | Fisher Scientific | Cat# 15340168 |
| **Spark 20M Multimode Microplate Reader** | Tecan | N/A |
| **NanoQuant Infinite M200** | Tecan | N/A |
| **Guava® easyCyte Benchtop Flow Cytometer** | Millipore | Guava easyCyte 6 2L |
| **MESO QuickPlex instrument** | Mesoscale Discovery Inc. | SQ 120 |
| **gentleMACS™ Dissociator** | Miltenyi Biotec | Cat# 130-093-235 |
| **Transwell Inserts (6.5 mm, 0.4 µm pore size)** | Corning | Cat# 3470 |
| **Cell Scraper** | Millipore Sigma | Cat# C5981-100EA |
| **Millicell EZ Slide 8-Well Chamber** | Millipore Sigma | Cat# PEZGS0816 |
| **Trypan Blue Stain** | Invitrogen | Cat# T10282 |
| **70 µm Cell Strainer** | Thermo Fisher Scientific | Cat# 22-363-548 |
| **Noyes Spring Scissors - Angled** | Fine Science Tools | Cat# 15013-12 |
| **16% Paraformaldehyde** | Electron Microscopy Sciences | Cat#15710 |
| **ProLong Glass Antifade Mountant** | Thermo Fisher Scientific | Cat#P36984 |
| **2.5% Normal Goat Serum** | Vector Laboratories | Cat#S-1012-50 |
| **2.5% Normal Horse Serum** | Vector Laboratories | Cat#S-2012-50 |

### Resource Availability

#### Lead Contact

Further information and requests for resources and reagents should be directed to and will be fulfilled by the lead contact, Pradipta Ghosh.

#### Materials Availability

This study has generated CRC organoid biobank, RNA and DNA from the organoids. These materials are available from the lead contact with a completed Materials Transfer Agreement and patented technology agreement following the guidelines of the University of California, San Diego.

#### Data and Code Availability

All transcriptomic datasets generated in this work has been deposited at NCBI GEO [GSE237623, GSE237624, GSE237625]. The data underlying all the figures and tables are available in the article and its online supplemental materials. The Boolean analysis framework is available at https://github.com/sinha7290/Prodiff.

## Method Details

### Computational Approaches

#### Data Processing

We built a human colon tissue database (n = 1,911) derived from the previously published “Human Colon Global Database,” refined by filtering for EpCAM and Albumin expression and restricted to the Affymetrix Human U133 Plus 2.0 platform (<u>GPL570</u>). The dataset was obtained from the NCBI Gene Expression Omnibus (GEO) repository^133–135^, as described previously^52^, and was expanded by including 68 additional adenoma samples and FACS-purified human colon crypt samples (<u>GSE31255</u>). This dataset contained experiments from 23 independent NCBI-GEO data-series (GSEs) from the same platform (<u>GPL570</u>). A list of the 23 NCBI-GEO GSEs contained within the “Human colon tissue database” is provided in (**Supplemental Information 1**). Then, the dataset was prepared for Boolean analysis by selecting genes that had the best dynamic range (identified by the biggest percentile range between the 10^th^ and 90^th^ percentile of low and high expression value based on the *StepMiner*^82^ threshold for each gene in this dataset). All training and validation dataset (**Supplemental Information 1**) were downloaded from the NCBI GEO website^133–135^. All gene expression datasets (**Supplemental Information 1**) were processed separately using the Hegemon data analysis framework^40,51,52^. We did not combine datasets that belong to two different platforms.

#### Boolean Analysis

*Boolean logic* is a simple mathematical relationship of two values, i.e., high/low, 1/0, or positive/negative. The Boolean analysis of gene expression data requires first the conversion of expression levels into two possible values. Here, *StepMiner*^82^ algorithm is used to perform Boolean analysis of gene expression data^7^. StepMiner is an algorithm designed to detect stepwise transitions in time-series gene expression data^82^. It fits step functions to expression profiles by identifying the sharpest changes in signal, corresponding to gene expression switching events. The algorithm evaluates all possible step positions and calculates the average expression on either side of each step to define constant segments. An adaptive regression approach is then used to select the step position that minimizes the sum of squared errors between the observed and fitted data. The selected step is used as the *StepMiner* threshold*. This* threshold is used to convert gene expression values into Boolean values. A noise margin of 2-fold change is applied around the threshold to determine intermediate values, and these values are ignored during Boolean analysis. Finally, a regression test statistic is computed to assess the significance of the identified step transition as follows:

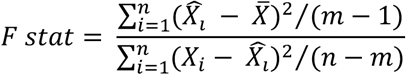

Where *X_i_* for *i* = 1 to *n* are the values, 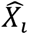 for *i* = 1 to *n* are fitted values. m is the degrees of freedom used for the adaptive regression analysis. 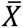 is the average of all the values: 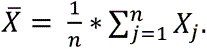 For a step position at k, the fitted values 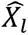 are computed by using 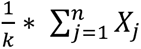 for *i* = 1 to *k* and 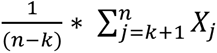 for *i* = *k* + 1 to *n*.

In a scatter plot, there are four possible quadrants based on Boolean values: (low, low), (low, high), (high, low), (high, high). In all our experimental settings, we applied the same logic to classify samples into CDX2-high and CDX2-low groups based on the dynamic range of CDX2 expression levels.

##### Invariant Boolean implication relationships

A Boolean implication relationship is observed if any one of the four possible quadrants or two diagonally opposite quadrants are sparsely populated. Based on this rule, there are six different kinds of Boolean implication relationships. Two of them are symmetric: equivalent (corresponding to the highly positively correlated genes), opposite (corresponding to the highly negatively correlated genes). Four of the Boolean relationships are asymmetric, and each corresponds to one sparse quadrant: (low => low), (high => low), (low => high), (high => high). BooleanNet statistics (Equations listed below) is used to assess the sparsity of a quadrant and the significance of the Boolean implication relationships ^32,50^. Given a pair of genes A and B, four quadrants are identified by using the *StepMiner* thresholds on A and B by ignoring the Intermediate values defined by the noise margin of 2-fold change (+/- 0.5 around *StepMiner* threshold). Number of samples in each quadrant are defined as a_00_, a_01_, a_10_, and a_11_. Total number of samples where gene expression values for A and B are low is computed using following equations.

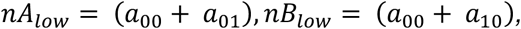

Total number of samples considered is computed using following equation.

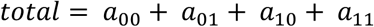

Expected number of samples in each quadrant is computed by assuming independence between A and B. For example, expected number of samples in the bottom left quadrant 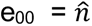 is computed as probability of A low ((a_00_ + a_01_)/total) multiplied by probability of B low ((a_00_ + a_10_)/total) multiplied by total number of samples.

Following equation is used to compute the expected number of samples for the low-low quadrant.

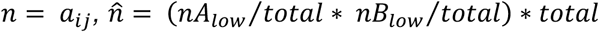

To check whether a quadrant (here low-low) is sparse, a statistical test for (e_00_ > a_00_) or 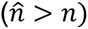 is performed by computing S_00_ and p_00_ using following equations. A quadrant is considered sparse if S_00_ is high 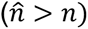 and p_00_ is small.

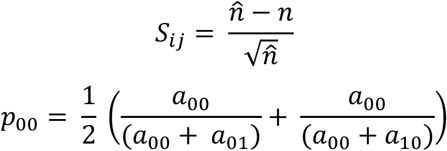

A threshold of S_00_ > sthr and p_00_ < pthr to check sparse quadrant. A Boolean implication relationship is identified when a sparse quadrant is discovered using following equation.

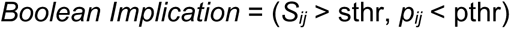

A relationship is called Boolean equivalent if top-left and bottom-right quadrants are sparse.

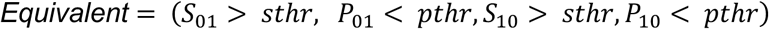

Boolean opposite relationships have sparse top-right (a_11_) and bottom-left (a_00_) quadrants.

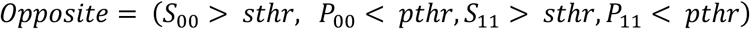

Boolean equivalent and opposite are symmetric relationship because the relationship from A to B is same as from B to A. Asymmetric relationship forms when there is only one quadrant sparse (A low => B low: top-left; A low => B high: bottom-left; A high=> B high: bottom-right; A high => B low: top-right). These relationships are asymmetric because the relationship from A to B is different from B to A. For example, A low => B low and B low => A low are two different relationships.

A low => B high is discovered if bottom-left (a_00_) quadrant is sparse and this relationship satisfies following conditions.

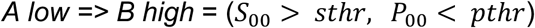

Similarly, A low => B low is identified if top-left (a_01_) quadrant is sparse.

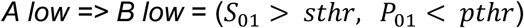

A high => B high Boolean implication is established if bottom-right (a_10_) quadrant is sparse as described below.

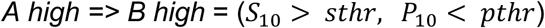

Boolean implication A high => B low is found if top-right (a_11_) quadrant is sparse using following equation.

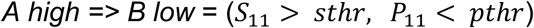

For each quadrant, a statistic S_ij_ and an error rate p_ij_ is computed. S_ij_ > 10 and p_ij_ < 0.15 are the thresholds used on the BooleanNet statistics to identify Boolean implication relationships (BIRs). False discovery rate is computed by randomly shuffling each gene and computing the ratio of the number of Boolean implication relationship discovered in the randomized dataset and original dataset. The false discovery rate for our dataset was less than 0.001.

Boolean analysis on the large human colon tissue database (n = 1911; Global 2018 GPL570 Colon Purged), uses a threshold of sThr = 10 and pThr = 0.15. We increased sThr and relax the pThr to focus on reasonable number of gene candidates. Boolean Implication analysis looks for invariant relationship across all the different types of samples regardless of the conditions and treatment protocols. Therefore, it does not distinguish the sample types when discovering Boolean implication relationships. We assume that there are fundamental invariant Boolean implication formula that are satisfied by every sample regardless of their type.

##### CANDiT (Cancer Associated Nodes for Differentiation Targeting)

We developed a machine learning framework, *CANDiT*, designed to identify actionable nodes that drive differentiation within cancer networks to guide therapeutic intervention. *CANDiT* is agnostic to tissue of origin and begins with a single ‘seed’ gene that meets two essential criteria: (i) it must be a lineage-defining transcriptional determinant along the stemness-to-differentiation axis in the cancer’s tissue of origin, and (ii) it must hold prognostic relevance in human tissue datasets, linking its expression to patient outcomes. Thus, the seed gene must bridge two levels of biological scale—cellular differentiation and clinical prognosis.

*CANDiT* constructs a Boolean implication network (BIN) by identifying all statistically significant pairwise Boolean implication relationships (BIRs)^8^ within a curated expression compendium, the Human Colon Global Database, as previously described^52^. The resulting BIN is a directed graph in which nodes represent genes and edges represent one of six possible Boolean relationships (e.g., high ⇒ high, high ⇒ low, low ⇒ high, low ⇒ low, Equivalent, Opposite). While Equivalence relationships often form a scale-free network, other asymmetric BIRs do not.

Boolean analysis was performed on genes with sufficient dynamic range to permit clear binary classification (high/low) via the *StepMiner* algorithm. Based on over a decade of experience with Boolean implication methods, BIRs are most reliable in datasets with >200 samples^136^. Our dataset was prepared for Boolean analysis by filtering genes that had a reasonable dynamic range of expression values. Genes with fewer than 5% of samples in either the high or low state were excluded to reduce noise and improve robustness. This filtering step ensures the resulting network highlights biologically meaningful and robust relationships.

##### Generation of Clustered Boolean Implication network

Clustering was performed in the Boolean implication network to dramatically reduce the complexity of the network. A clustered Boolean implication network (CBIN) was created by clustering nodes in the original BIN by following the equivalent BIRs. One approach is to build connected components in an undirected graph of Boolean equivalences. However, because of noise, the connected components become internally inconsistent e.g., two genes opposite to each other become part of the same connected component. In addition, the size of clusters became unusually big with almost everything in one cluster. To avoid such a situation, we need to break the component by removing the weak links. To identify the weakest links, we first computed a minimum spanning tree for the graph and computed the Jaccard similarity coefficient for every edge in this tree. Ideally if two members are part of the same cluster, they should share as many connections as possible. A threshold of 0.7 is considered for the Jaccard similarity coefficient and if they share less than 70% of their total individual connections (Jaccard similarity coefficient less than 0.7) the edges are dropped from further analysis. Thus, many weak equivalences were dropped using the above algorithm leaving the clusters internally consistent. We removed all edges that have Jaccard similarity coefficient less than the selected threshold and built the connected components with the rest. The connected components were used to cluster the BIN which is converted to the nodes of the CBIN. The choice of the threshold on the Jaccard similarity coefficient play an important role in determining the size and the number of clusters as well as whether they are internally consistent. A new graph was built that connected the individual clusters to each other using Boolean relationships. The link between two clusters (A, B) was established by using the top representative node from A that was connected to most of the members of A and sampling 6 nodes from cluster B and identifying the overwhelming majority of BIRs between the nodes from each cluster.

Here, CBIN was created using our pooled dataset (human colon tissue database (n = 1911) to capture the differentiation events in colon tissue (**Figure S1C**). The edges between the clusters represented the Boolean relationships that are color-coded as follows: orange for low => high, dark blue for low => low, green for high => high, red for high => low, light blue for the equivalent and black for the opposite. A subnetwork is selected using low=>low (blue), high => low (red) and opposite (merged with high=>low as red) edges among the top 10 clusters (**Figure S1D**).

##### Charting Boolean paths

Boolean paths have been previously leveraged to predict the underlying time series events in biological processes such as B cell differentiation^32,33^ and early differentiation events in cancer stem cell^40,43,51,52^. The core algorithm enabling this, *MiDReG* (Mining Developmentally Regulated Genes), uses two seed genes infer intermediate genes and transitional states within a developmental or differentiation process. *MiDReG* identifies these intermediates through a series of asymmetric BIRs ^10^.

In this study, we apply the *MiDReg* framework to traverse the clustered Boolean Implication network, identifying gene clusters whose start and end points may define biologically meaningful differentiation trajectories in colon tissue. The asymmetric BIRs offer a directional, causal logic that distinguishes this network from conventional co-expression or correlation-based gene networks.

A Boolean path refers to a directed sequence of BIR-linked nodes (genes) within the BIN. A simple Boolean path includes two nodes and a single directed edge, while a complex Boolean path involves multiple nodes and edges, representing multi-step transitions between regulatory states. These paths allow for a high-resolution mapping of potential developmental hierarchies or transitions embedded in steady-state transcriptomic data.

##### Ordering samples based on composite scores along Boolean paths

Each Boolean path comprises one or more clusters. To assign a composite score to each sample, we first computed a normalized average expression value for the genes within each cluster. Normalization was performed using a modified Z-score centered around the *StepMiner* threshold (SThr), defined as: (formula = (expr - SThr)/3*stddev).

Next, a weighted linear combination of these cluster averages was used to generate a final composite score for each sample. Weights along the Boolean path were assigned to either monotonically increase or decrease—ensuring that the sample ordering remained consistent with the logical sequence dictated by the asymmetric BIRs. Clusters highly expressed in disease samples were assigned a weight of +1, while those predominant in healthy samples received a weight of -1. The directionality of the Boolean path—from healthy to disease states—guided this weighting scheme. Samples were then ordered based on their final composite score, enabling biologically meaningful stratification.

The genes contained within each cluster of the Boolean network are detailed in **Supplemental Information 2**.

###### Measurement of Classification Strength or Prediction Accuracy

To evaluate classification strength and prediction accuracy, Receiver Operating Characteristic (ROC) curves were generated for each gene. These curves assess the performance of a binary classifier, e.g., high vs. low *StepMiner* normalized gene expression levels, across varying discrimination thresholds. ROC curves plot the True Positive Rate (TPR) against the False Positive Rate (FPR) at multiple threshold levels. The Area Under the Curve (AUC) quantifies the classifier’s ability to correctly distinguish between the various types of samples, in this instance, CRC or healthy and treated or not with drug. ROC-AUC values were computed using the Python Scikit-learn package.

###### Correlation Coefficients Analysis

Linear correlation analysis was performed between ΔCT values of *CCDC88A*, CDX2 and *PRKAB1* of all the CRC PDOs with their IC50 values. Regression analysis was performed using SciPy Stat package. Scatter plot with regression lines were generated using a Seaborn package (0.12.1).

###### Integrated Differential Expression Analysis

Differential gene expression analysis was performed using RNAseq raw count data processed through DESeq2^137^. Genes with an absolute log2 fold change [FC] > 10 and an adjusted p-value <0.05 were considered differentially expressed genes (DEGs). Pathway analysis of gene lists was carried out via the Reactome database and algorithm ^138^. Reactome identifies signalling and metabolic molecules and organizes their relations into biological pathways and processes. A complete catalogue of DEGs and reactome pathway enrichments are provided in **Supplemental Information 4** and **Supplemental Information 5**, respectively. The 50-gene signature that emerged from the integration of all DEGs is provided in **Supplemental Information 6**.

###### Survival Analysis

*StepMiner* based survival analysis was performed by dividing the patients into two groups, based on high or low levels of expression of the 50-genes signature of therapeutic response [composite score expression values], as determined by implementing the *StepMiner*-derived threshold for each cohort (https://github.com/sinha7290/Prodiff). Log-rank analysis and visualization were performed using GraphPad Prism version 9.1. Default settings of GraphPad Prism ignore events at time zero (t0) for the assessment of the impact of treatment modalities on outcome (https://www.graphpad.com/support/faq/events-deaths-at-time-zero-in-survival-analysis/). These default settings were manually modified to include all samples (without exception) in the cohort for which outcome was known. Survival analysis on KRAS^WT^ patient groups based on microsatellite stability status was performed using KM-plotter^129^ based on *auto selected best percentile cutoff* of 50 genes.

###### Univariate and Multivariate Analyses

To assess which factor(s) may influence the IC50 of the PDOs, multivariate regression has been performed over all the clinical conditions. Here, the statsmodels module from python has been used to perform Ordinary least-squares (OLS) regression analysis of each of the variables. The p-value for each term tests the null hypothesis that the coefficient is equal to zero (no effect).

We also performed univariate analysis on GSE39582 using the same OLS regression analysis to assess the impact of each variables individually, such as tumor stage, age, gender, tumor location, CIMP status, mutation status (*BRAF* and *TP53*) and gene expression [either *CDX2* alone or the composite score of the 50-gene signature of therapeutic response, both converted into a binary value, high vs low, as determined by *StepMiner* threshold] on survival. The analysis was done on the entire cohort, as well as on *KRAS* WT and *KRAS* mutant patients as two independent sub-cohorts. Both the univariate and multivariate analysis data are represented based on the coefficient values of associated variables and their upper and lower bounds of 95% confidence interval as the error bars using Matplotlib. The significance of the variables was determined by t-test; * = p < 0.05, ** = p < 0.01, and *** = p < 0.001.

###### Unsupervised Clustering and Heatmap

Expression patterns of the genes that are differentially expressed in *CDX2*-low and -high groups, with or without PF treatment, are clustered without bias based on their z-normalized cpm expression values, in all the samples. The data is visualized using the seaborn clustermap package (v 0.12) in python.

###### Bulk RNAseq Deconvolution

*In silico* deconvolution of bulk RNA-seq data from xenograft models was performed using the Granulator R package^139^ to estimate monocyte abundance. Cell-type abundance estimates were normalized using the immune cell signature matrix developed by Monaco et al^140^.

###### Statistical Tests

Optimal gene expression cut-off values were determined using the *StepMiner* algorithm within each individual dataset^82^. Gene signatures were used to classify sample categories, and multi-class classification performance was evaluated using ROC-AUC (receiver operating characteristic area under the curve) values. Statistical comparisons were performed using Welch’s two-sample t-test (unpaired, unequal variance and sample size) via the scipy.stats.ttest_ind function in Python (version 0.19.0; equal_var=False). Multiple hypothesis correction was applied using the Benjamini-Hochberg method (fdr_bh) via statsmodels.stats.multitest.multipletests. Differential expression analysis was conducted using DESeq2 in R (version 1.16.1). Kaplan–Meier survival analysis and log-rank tests for statistical significance were performed using GraphPad Prism 9.1. Violin and bubble plots were generated using Seaborn (Python, version 0.12). Boolean implication relationships between genes were determined using the BooleanNet algorithm (criteria: statistic > 3 and error rate < 0.1). Power calculations estimated effect sizes (mean difference divided by standard deviation) of 2.5 in PDOs and 1.0 in animal experiments for CDX2 induction using the PRKAB1 agonist. A minimum of 4 PDOs and 17 animals per group was determined to provide adequate power (α = 0.05, power = 0.80).

### Experimental Approaches

#### Chemical Synthesis of PRKAB1 Agonist

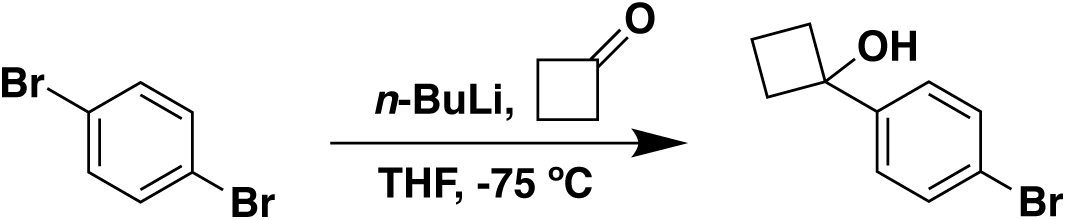

##### 1-(4-Bromophenyl)cyclobutan-1-ol

An oven-dried 50 mL round-bottom flask was charged with 1,4-dibromobenzene (3.43 g, 14.6 mmol, 1.2 equiv) and anhydrous tetrahydrofuran (THF, 14 mL) and was cooled in a dry ice-acetone bath (−75 °C). *n*-Butyllithium (*n*-BuLi, 2.5 M in hexane, 5.8 mL, 14.6 mmol, 1.2 equiv) was added dropwise. After stirring for 30 min., cyclobutanone (0.850 g, 12.1 mmol, 1 equiv) was added dropwise. The reaction mixture was stirred for 60 min at -75°C. The reaction was quenched by the addition of a saturated NH_4_Cl (aq) solution (∼7 mL). Water was added (∼30 mL) and the solution mixture was extracted twice with Ethyl Acetate (EtOAc, ∼100 mL x 2), the combined organics were washed with saturated brine (∼50 mL), dried over MgSO4, filtered, and concentrated *in vacuo*. The product was purified via SiO2 column chromatography (using a gradient of 0% to 10% to 15% EtOAc in hexanes as eluent) to give the title compound as a white solid (2.40 g, **87% yield**). 1H NMR (300 MHz, CDCl3) δ (ppm) = 7.49-7.46 (d, 2H), 7.37-7.34 (d, 2H), 2.54-2.46 (m, 2H), 2.39-2.29 (m, 2H), 2.28 (s, 1H), 2.08-1.94 (m, 1H), 1.75-1.61 (m, 1H).

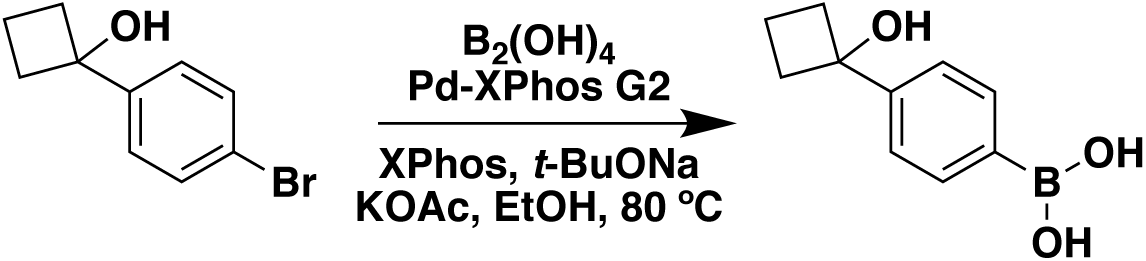

##### (4-(1-Hydroxycyclobutyl)phenyl)boronic acid

An oven-dried 100 mL round-bottom flask was charged with 1-(4-bromophenyl)cyclobutan-1-ol (2.00 g, 8.81 mmol, 1 equiv) and anhydrous ethanol (25 mL). Sodium tert-butoxide (8.5 mg, 0.088 mmol, 0.01 equiv), oven-dried potassium acetate (1.82 g, 18.5 mmol, 2.1 equiv), XPhos Pd G2 (chloro(2-dicyclohexylphosphino-2′,4′,6′-triisopropyl-1,1′-biphenyl)-[2-(2′-amino-1,1′-biphenyl)]palladium(II), 36 mg, 0.044 mmol, 0.005 equiv), and XPhos (2-dicyclohexylphosphino-2′,4′,6′-triisopropylbiphenyl, 42 mg, 0.088 mmol, 0.01 equiv) were added and the mixture was sparged with Argon. Tetrahydroxydiboron (1.58 g, 17.6 mmol, 2.0 equiv) was added and was stirred at 80 °C for 3 hr. The mixture was cooled to room temperature, and the solids were filtered through celite to give a light-yellow solution. Water was added (∼20 mL) and the solution mixture was extracted twice with EtOAc (∼50 mL x 2), the combined organics were washed with saturated brine (∼20 mL), dried over MgSO4, filtered, and concentrated *in vacuo*. The product was purified via SiO2 column chromatography (using a gradient of 50% to 60% to 70% EtOAc in hexanes as eluent) to give the title compound as a white solid (0.900 g, **53% yield**). 1H NMR (300 MHz, DMSO-d6) δ (ppm) = 7.96 (s, 2H), 7.77-7.74 (d, 2H), 7.45-7.43 (d, 2H), 5.45 (s, 1H), 2.42−2.33 (m, 2H), 2.30−2.24 (m, 2H), 1.98−1.84 (m, 1H), 1.71−1.56 (m, 1H).

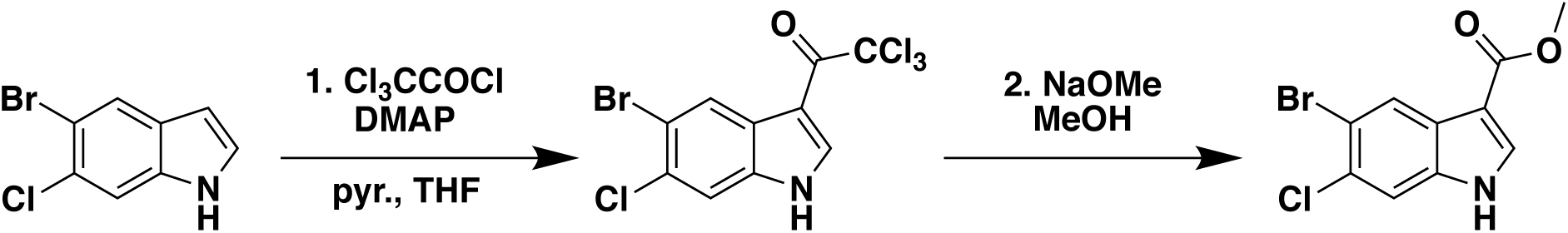

##### Methyl 5-Bromo-6-chloro-1H-indole-3-carboxylate

An oven-dried 100 mL round-bottom flask was charged with 5-bromo-6-chloro-1H-indole (2.10 g, 9.11 mmol, 1 equiv), DMAP (Dimethylaminopyridine, 111 mg, 0.911 mmol, 0.1 equiv), pyridine (1.87 g, 23.7 mmol, 2.6 equiv), and anhydrous THF (14 mL) and then cooled to 0 °C. Trichloroacetyl chloride (3.98 g, 21.9 mmol, 2.4 equiv) was then added dropwise, and then stirred for 3 days at room temperature. The mixture was then cooled to 0 °C, and methanol (3.7 mL, 10 equiv) was added dropwise, then 25% sodium methoxide in methanol solution (4.2 mL, 2 equiv) was added dropwise. The mixture was stirred at 55 °C for 2 hr. Water was added (∼20 mL) and the solution mixture was extracted twice with EtOAc (∼50 mL x 2). The combined organics were washed with saturated brine (∼20 mL), dried over MgSO4, filtered, and concentrated *in vacuo*. The product was purified via SiO2 column chromatography (using a gradient of 20% to 30% EtOAc in hexanes as eluent) to give the title compound as an off-white solid (1.67 g, **63% yield**). 1H NMR (300 MHz, CDCl_3_): δ (ppm) = 8.52 (br s, 1H), 8.45 (s, 1H), 7.92 (d, 1H), 7.56 (s, 1H), 3.93 (s, 3H).

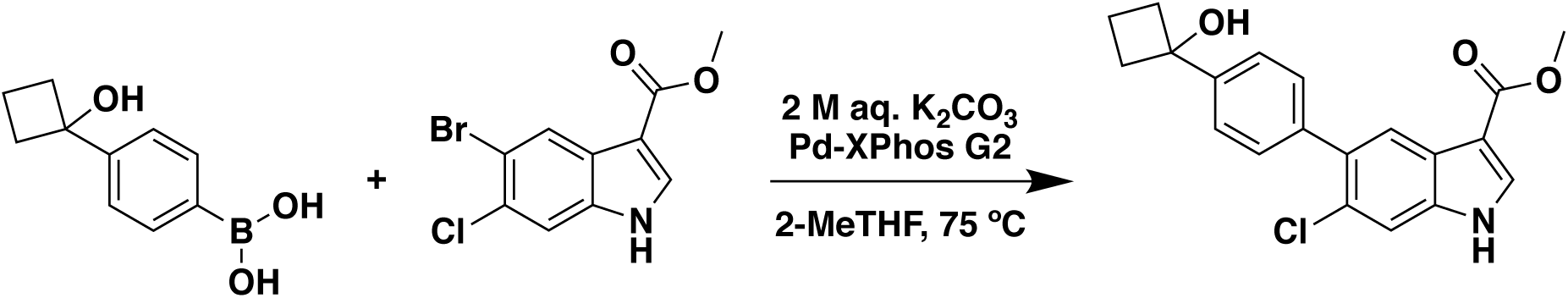

##### Methyl 6-Chloro-5-(4-(1-hydroxycyclobutyl)phenyl)-1H-indole-3-carboxylate

An oven-dried 10 mL round-bottom flask was charged with methyl 5-bromo-6-chloro-1H-indole-3-carboxylate (280 mg, 0.970 mmol, 1 equiv), (4-(1-hydroxycyclobutyl)phenyl)boronic acid (205 mg, 1.07 mmol, 1.1 equiv), 2-MeTHF (2 mL), XPhos Pd G2 (chloro(2-dicyclohexyl-phosphino-2′,4′,6′-triisopropyl-1,1′-biphenyl)-[2-(2′-amino-1,1′-biphenyl)] palladium(II), 16 mg, 0.019 mmol, 0.02 equiv). Aqueous K_2_CO_3_ (362 mg, 2.62 mmol, 2.7 equiv in 1.1mL water, ∼2.4 M) was added, and the mixture was sparged with Argon. The reaction mixture was heated to 75°C overnight (18 hrs). After cooling, the solvent was evaporated to dryness. Water was added (∼10 mL) and the solution mixture was extracted three times with EtOAc (∼10 mL x 3), the combined organics were washed with saturated brine (∼10 mL), dried over MgSO4, filtered, and concentrated *in vacuo*. The product was purified via SiO2 column chromatography (using a gradient of 30% to 40% to 50% EtOAc in hexanes as eluent) to give the title compound as a white solid (170 mg, **49% yield**). 1H NMR (300 MHz, DMSO-d6): δ (ppm) = 12.08 (br s, 1H), 8.17 (s, 1H), 7.93 (s, 1H), 7.66 (s, 1H), 7.58-7.55 (d, 2H), 7.41-7.39 (d, 2H), 5.55 (s, 1H), 3.79 (s, 3H), 2.48−2.41 (m, 2H), 2.35−2.26 (m, 2H), 2.00−1.91 (m, 1H), 1.76−1.64 (m, 1H).

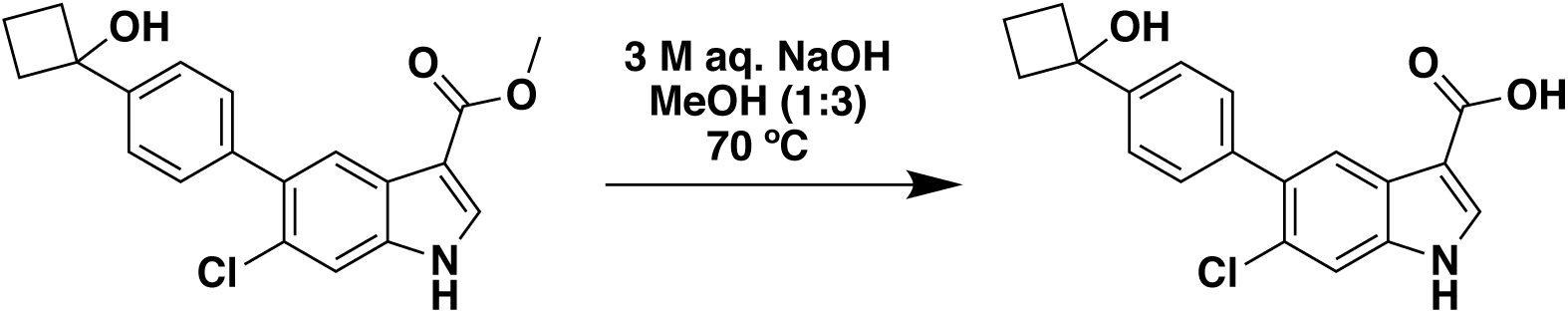

##### 6-Chloro-5-(4-(1-hydroxycyclobutyl)phenyl)-1H-indole-3-carboxylic acid (PF06409577)

A 4-dram vial under Argon was charged with methyl 6-chloro-5-(4-(1-hydroxycyclobutyl)phenyl)-1H-indole-3-carboxylate (140 mg, 0.393 mmol, 1 equiv), 3M NaOH (aq) solution (0.787 mL, 2.36 mmol, 6 equiv), and methanol (MeOH, 2.36 mL). The reaction mixture was heated to 70°C overnight (18 hrs). After cooling, the solvent was evaporated to dryness. Water was added (∼10 mL), the pH was adjusted to ∼2 using 1M HCl (aq) solution, and the solution mixture was extracted three times with EtOAc (∼10 mL x 3), the combined organics were washed with saturated brine (∼10 mL), dried over MgSO4, filtered, and concentrated *in vacuo*. The product was purified via SiO2 column chromatography (using a gradient of 30% to 60% Acetonitrile in Dichloromethane as eluent) to give the title compound as a white solid (54 mg, **40% yield**). 1H NMR (300 MHz, DMSO-d6): δ (ppm) = 11.94 (br s, 1H), 8.05 (s, 1H), 7.96 (s, 1H), 7.63 (s, 1H), 7.58-7.55 (d, 2H), 7.41-7.39 (d, 2H), 5.53 (s, 1H), 2.48−2.40 (m, 2H), 2.35−2.26 (m, 2H), 2.00−1.89 (m, 1H), 1.74−1.63 (m, 1H).

HPLC characterization was conducted using the Agilent 1260 Infinity II Quaternary Pump System with a 150 mm x 3 mm, 2.7 mm particle size C18 column (693975-302T). Solvent conditions (Solvent A: H2O (0.1% TFA), Solvent B: Acetonitrile): 0-2 min isocratic 5% B, 2-20 min gradient to 95% B, 20-22 min isocratic 95% B, at a flow rate of 1 mL/min. The chromatogram was monitored at 254 nm wavelength and the compound had a retention time of 11.956 min. The purity of this final product was 96% by HPLC.

#### Cell Culture

##### HCT-116

This human colon cancer cell line was obtained from the American Type Culture Collection (ATCC) and maintained in Roswell Park Memorial Institute 1640 (RPMI-1640) medium (Gibco-BRL). The medium was supplemented with 10% fetal calf serum (Hyclone), 1% L-Glutamine, and 1% penicillin/streptomycin (Gibco-BRL). The cells were incubated at 37°C in a 5% CO2 incubator and were routinely passaged at a dilution of 1:5 to 1:10.

##### SW480

This human colon cancer cell line was obtained from the ATCC and maintained in DMEM/F12, supplemented with 10% FBS. The cells were incubated at 37°C in a 5% CO2 incubator and were routinely passaged at a dilution of 1:2 to 1:5.

##### DLD1

This human colon cancer cell was obtained from the ATCC and were cultured using RMPI media containing 10% FBS. The cells were incubated at 37°C in a 5% CO2 incubator and were routinely passaged at a dilution of 1:5 to 1:10.

##### Caco2

This human colorectal cancer cell line was obtained from the ATCC and was cultured in high glucose DMEM/F12 supplemented with 10% FBS and 1% penicillin/streptomycin (Cytiva). The cells were incubated at 37°C in a 5% CO2 incubator and were routinely passaged at a dilution of 1:3 to 1:8.

#### MTT (3-[4,5-dimethylthiazol-2-yl]-2,5 diphenyl tetrazolium bromide) assays on CRC cell lines

Cell survival was measured using the MTT reagent and cells cultured in 96-well plates. HCT116 and Sw480 cells were cultured and treated with different concentrations of PF compound (0, 1, 5, 10, 20 µM) using DMSO as a negative control for 16-48 h in various assays. Then the cell lines were incubated with MTT for 4 hr at 37°C. After incubation, culture media was removed, and replaced with phosphate buffered saline (PBS), and 150 μl of DMSO was added in order to solubilize the MTT formazan crystals. Optical density was determined at 590 nm using a TECAN plate reader. At least three independent experiments were performed.

#### Cell Cycle and Apoptosis Analyses

Cell cycle distribution and apoptosis were assessed using the Guava Cell Cycle Reagent (Millipore Sigma) and the Annexin V/Propidium Iodide (PI) staining kit (Thermo Fisher Scientific), respectively, following the manufacturers’ protocols. Samples were analyzed on a BD LSR II flow cytometer, and data were processed using FlowJo software (FlowJo LLC, Ashland, OR).

#### Cell Toxicity Assays

Confluent flasks of HCT116 and Caco2 cells were rinsed with PBS and detached using 1X trypsin-EDTA in PBS. Cells were resuspended in 1 mL of their respective medias (HCT116: McCoy’s 5A modified medium with 10% FBS and 1% Pen/Strep; Caco2: High glucose DMEM with 10% FBS, 1% NEAA, 1% P/S). Prior to seeding, 50 µL of media was added to each well of an Agilent xCELLigence 96-well impedance plate, and baseline cell index readings (2 sweeps) were recorded using the Agilent eSight live-cell analysis system. Cells were then counted and seeded at 10,000 cells/well in 50 µL of media. Cell index measurements were collected at 15-minute intervals for 24 hours to monitor attachment and proliferation.

After 24 hours, the assay was paused to allow treatment with compounds or vehicle. Each well received 100 µL of treatment media containing 2X concentrations of the PF compound or DMSO (vehicle), along with 2X Annexin V. The plate was returned to the instrument and continuous monitoring resumed. Cell index measurements and live-cell images (4/well) were collected throughout the remainder of the 126-hour assay. Cell index values were normalized to the final pre-treatment time point. Dose-response curves and Annexin-based toxicity were assessed at the 126-hour endpoint.

#### PDO Toxicity Assay

PDOs were cultured for 72 hours prior to treatment. Organoids were then exposed to complete medium containing either PF compound (5 μM or 20 μM), DMSO (vehicle), or hydrogen peroxide (H₂O₂, positive control), with 500 nM eTox Red added to detect cell death. After 72 hours of treatment, Hoechst 33342 (5 μg/mL) was added to label nuclei, followed by a PBS wash and replacement with phenol red- and serum-free DMEM. Organoids were imaged using a BioTek Cytation 10 system. eTox Red-positive cells (defined by red fluorescence within nuclei) were quantified using BioTek Gen5 software, and the percentage of positive cells was calculated per condition. Data were analyzed and visualized using GraphPad Prism.

#### Animal Model

Athymic nude male mice ages 4–6 weeks purchased from Jackson Laboratories (Bar Harbor, ME) were utilized for this study. Mice were maintained in a barrier facility with high-efficiency particulate air filtration and fed an autoclaved laboratory diet. Prior to surgical procedures, mice were anesthetized with an intraperitoneal injection of ketamine and xylazine reconstituted in phosphate-buffered saline (PBS). When the study concluded or if tumor burden became too large, defined as tumor volume > 1500 cm^3^, mice were euthanized with CO2 inhalation followed by cervical dislocation. This study was carried out in strict accordance with the recommendations in the Guide for the Care and Use of Laboratory Animals of the National Institutes of Health. All animal studies were approved by the San Diego Veterans Administration Medical Center Institutional Animal Care and Use Committee (protocol A17-020).

##### Tumor establishment with cell injection

Subcutaneous injection of HCT-116 cells (1 x 10^6^) reconstituted in PBS and Matrigel Matrix (Corning, NY) was performed on the bilateral shoulders and flanks of nude mice. Tumors were allowed to grow until 50 mm^3^ in diameter. The tumors were then resected and divided into 1 mm^3^ pieces for subcutaneous implantation.

##### Tumor establishment with tumor implantation

Subcutaneous models were established by surgical implantation of 1 mm^3^ HCT-116 tumor fragment into the bilateral flank of nude mice. A small incision was made on the back of the mice and tumor fragment was directly implanted in the bilateral flank. The skin was closed with 6-0 nylon suture (Ethicon Inc., Sommerville. NJ). Tumors were allowed to grow until 150 mm^3^ in diameter.

#### Immunohistochemistry

Tumor sections of 4 μm thickness were cut and placed on glass slides coated with poly-L-lysine, followed by deparaffinization and hydration. Heat-induced epitope retrieval was performed using sodium citrate buffer (pH 6) or Tris-EDTA (pH9) in a pressure cooker. Tumor sections were incubated with 3% hydrogen peroxide for 5-15 min to block endogenous peroxidase activity, followed by incubation with primary antibody overnight in a humidified chamber at 4°C. Antibodies used for immunostaining were anti-CDX2 (1:300, rabbit host, cat no. ab76541), AMPK β1 (1:200, rabbit host, cat no. 10308-1-AP), anti-pSer^245^GIV (1:50, anti-rabbit), anti-YAP (clone 63.7; Santa Cruz: sc-101199), anti-phospho(p)YAP Ser^127^ (Cell Signaling Technology Cat#13008S) and anti-β-Catenin (1:300, mouse host, SC-7963). Antigen retrieval protocols were optimized based on manufacturer’s protocols. Immunostaining was visualized with a labelled streptavidin–biotin using 3,3′-diaminobenzidine as a chromogen and counterstained with hematoxylin. Immunohistochemistry (IHC) images were randomly sampled at different regions of interest (ROI). The ROIs were analysed using IHC Profiler^141^. IHC Profiler uses a spectral deconvolution method of DAB/hematoxylin colour spectra using optimized optical density vectors of the colour deconvolution plugin to properly segregate the DAB colour spectra. The histogram of the DAB intensity was divided into 4 zones: high positive (0–60), positive (61–120), low positive (121–180), and negative (181–235). High positive, positive, and low positive percentages were combined to compute the final percentage positive for each ROI. The range of values for the percent positive was compared among the different experimental groups.

#### Immunofluorescence

Tumor sections of 4 μm thickness were cut and placed on glass slides coated with poly-L-lysine, followed by deparaffinization and hydration. Heat-induced epitope retrieval was performed using Tris-EDTA (pH9) in a pressure cooker. Samples were blocked for 1 hr using an in-house blocking buffer (1% BSA and 0.01% Tween-20 in TBS). Primary antibodies were diluted in blocking buffer and allowed to incubate overnight at 4°C; antibodies used were E Cadherin (H-108) cat no. SC7870, 1:50, and Anti-Beta Catenin cat no. SC7963, 1:50. Secondary antibodies, Goat anti-mouse IgG H&L secondary antibody, Alexa Fluor 488 and Goat anti-rabbit IgG H&L secondary antibody, Alexa Fluor 594, were diluted in blocking buffer and allowed to incubate for 2 hr in the dark. ProLong Glass was used as a mounting medium. #1 Thick Coverslips were applied to slides and sealed. Samples were stored at 4°C until imaged.

#### Quantification of Nuclear Area

Formalin-fixed paraffin-embedded (FFPE) xenograft sections were stained with hematoxylin and eosin (H&E) using standard protocols. Whole slide images were acquired using a high-resolution slide scanner. Representative fields from vehicle- and PF-treated xenografts were exported as individual TIFF files for image analysis. To quantify nuclear morphology, images were processed using ImageJ with the StarDist plugin (v0.3.0), trained on an H&E-compatible model for nuclear detection. Each image was converted to 8-bit grayscale, and contrast was adjusted as needed. StarDist-generated nuclear segmentation masks were used to define regions of interest, and nuclear area was quantified for all detected nuclei in each field. Data visualization and statistical analysis were performed using GraphPad Prism.

#### Patient-derived Organoid (PDO) Culture

Twenty-three colorectal carcinoma (CRC) human patient-derived model (PDM) organoids were acquired from the Human Cancer Models Initiative (HCMI) catalogue of the American Type Culture Collection (ATCC). Five healthy colon PDOs were used from the biobank of the UC San Diego HUMANOID Center of Research Excellence (CoRE) at the University of California, San Diego, USA^142^. Organoids were cultured and passaged according to previously described methods^79,142–147^. For maintenance, organoids were cultured in Matrigel in 12- or 24-well tissue culture plates incubated at 37°C, 5% CO_2_. Organoids were provided 50% L-WRN-conditioned medium supplemented with differentiation inhibitors (Intestigrow; Cat#: HUM2019-125, HUMANOID™, San Diego, CA). Organoids were subcultured every 7-14 days, dissociated by non-enzymatic digestion in TrypLE and passaged to achieve the desired density. All relevant demographic, clinical and tumor mutation related information is provided in **Supplemental Information 3**.

#### Estimation of IC_50_ by MTT Assays on PDOs

To attain IC50 values for CRC PDM lines and healthy colon organoid lines in response to PF-06409577, thiazolyl blue tetrazolium bromide (MTT) reduction assays were performed, based on a previous protocol ^148^ with modifications. Prior to experiment, 5×10^3^-1×10^4^ dissociated single cells were seeded in Matrigel in 96-well tissue culture plates. After 3 days in culture, organoid images were captured on EVOS XL Core Imaging System. Organoids were administered complete medium or complete medium treated on alternate days with 1 µM, 5 µM, 10 µM, 20 µM, or 40 µM PF-06409577 (final concentration), while ensuring that only 50% of the media was replenished during subsequent repeated treatments. After 7-9 days of treatment, when apoptosis was most apparent, organoids were imaged and MTT reagent was added to each well for a final MTT concentration of 588 µg/ml. Organoids were incubated in MTT for 4 hours at 37°C, 5% CO_2_, after which images were acquired to document the extent of formazan formation. To prepare the plate for quantification, Matrigel was dissolved in 2% SDS, followed by formazan dissolution in dimethyl sulfoxide. The optical density of each well was measured at 562 nm using Tecan Spark Multimode Microplate Reader^148^. The measured optical densities at each concentration were used to calculate an IC50 for each CRC PDO and healthy colon line.

#### Immunoblotting

To verify the expression of markers of stemness and differentiation, equal aliquots of whole-cell lysates (prepared using RIPA buffer) were loaded on a 8% SDS PAGE gel, and immunoblotting was carried out for various targets as described previously^149^. Immunoblots were analyzed using a Gel Doc system (BioRad, Hercules, CA).

#### Flow Cytometry

To produce single-cell suspension from CRC PDOs for fluorescence-activated single cell sorting (FACS), dissociated single cells were seeded in Matrigel in 24-well tissue culture plates. After 3 days in culture, organoids were administered complete medium or complete medium treated with 1 µM, 5 µM, 10 µM, 20 µM, or 40 µM PF-06409577. After 7-9 days of treatment, organoids were collected in PBS-EDTA. Collected organoids were dissociated in TrypLE for subsequent analysis by flow. Apoptotic cell quantification was performed using the annexin V/propidium iodide (PI) staining kit (Thermo Fisher Scientific), according to the manufacturer’s instructions. Cells were quantified on a BD LSR II flow cytometer and analyzed using FlowJo software (FlowJo, Ashland, OR, USA).

#### RNA Extraction Transcriptomic Studies

To collect RNA from CRC PDOs, dissociated single cells were seeded in Matrigel in 24-well tissue culture plates. After 3 days in culture, organoids were administered complete medium or complete medium treated with 10 µM PF-06409577. Prior to treatment with PF-06409577 and after 2 days of treatment, whole organoids were collected in Dulbecco’s phosphate-buffered saline with 0.5 mM ethylenediaminetetraacetic acid (PBS-EDTA). Organoids were subsequently incubated in cell recovery solution, pelleted, and stored at -80°C. Total RNA was extracted using the Quick-RNA Microprep Kit and concentration was measured using Nanodrop One^C^ Spectrophotometer. For qPCR, first-strand cDNA synthesis was performed using qScript cDNA SuperMix.

For RNA sequencing, isolated RNA has been processed for RNA sequencing in the Illumina NovaSeq 6000 platform. Fastq sequence files have been mapped using the human GRCh38 genome. Log normalized CPM expression files are submitted to NCBI GEO [GSE237623, GSE237624, GSE237625].

#### Ethics Statement

All animal studies were conducted in strict accordance with the NIH *Guide for the Care and Use of Laboratory Animals* and were approved by the Institutional Animal Care and Use Committee of the San Diego Veterans Administration Medical Center (protocol A17-020; PI Bouvet). Colorectal cancer patient-derived organoids (PDOs) were obtained from the American Type Culture Collection (ATCC), a nonprofit global biorepository and distributor of organoid models curated under the Human Cancer Models Initiative (HCMI) to enhance reproducibility. Deidentified case-associated data, including molecular characterizations and tissue origins, were used in this study; harmonized datasets are publicly accessible via the NCI’s Genomic Data Commons. Healthy colon PDOs were derived from individuals undergoing routine colonoscopy for cancer screening at UC San Diego, with informed consent under an IRB-approved protocol (#190105; PI Ghosh).

