## Supplementary material for "Machine Learning–Guided Differentiation Therapy Targets Cancer Stem Cells in Colorectal Cancers": SOM

### INVENTORY OF SUPPLEMENTARY MATERIALS

- Supplemental Figure and Legends (Figure S1-S8)
- Supplemental Tables (Uploaded separately as Excel spreadsheets; Table S1-S7)

### SUPPLEMENTARY FIGURES AND LEGENDS

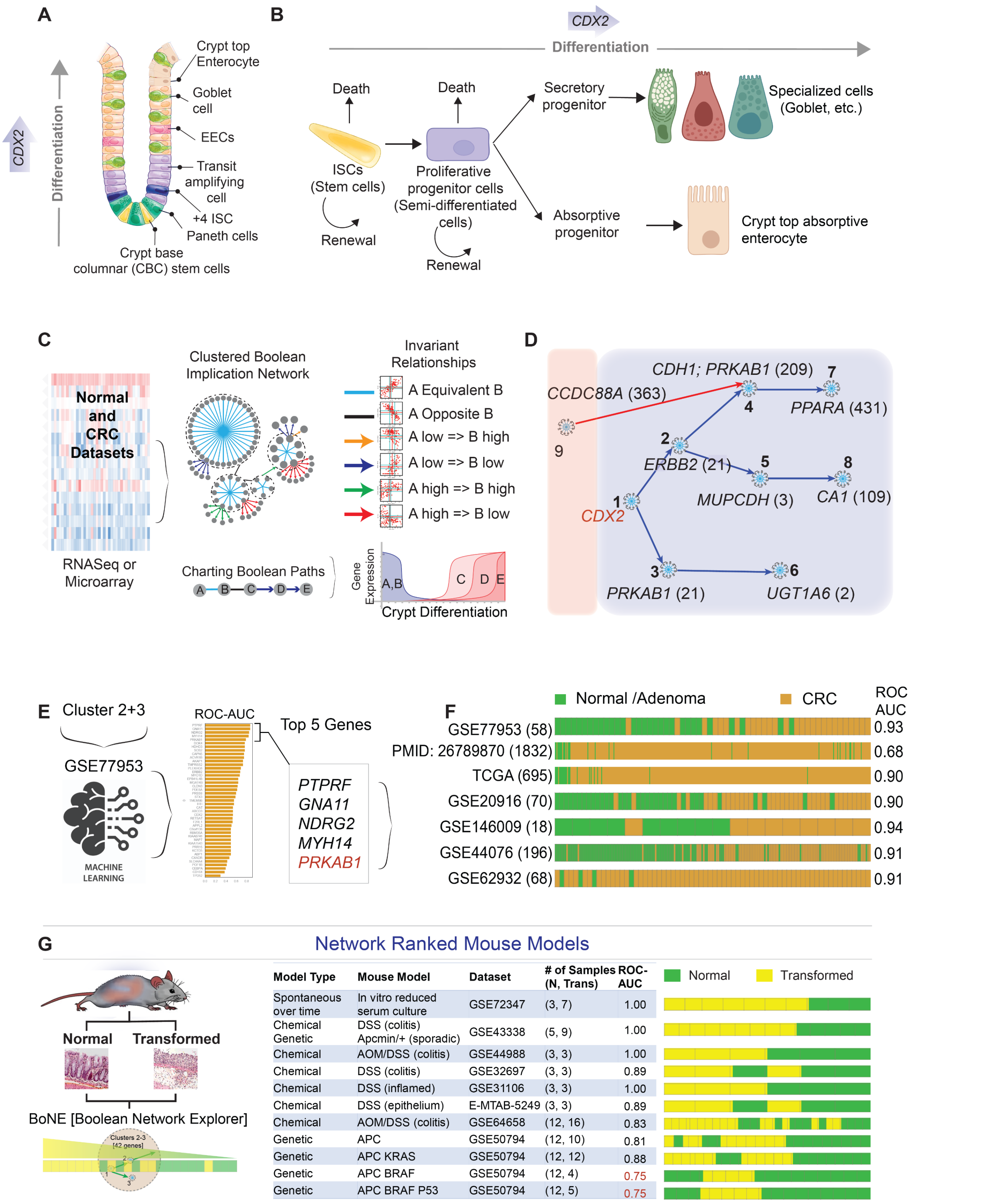


**Figure S1** [Related to Figure 1].

**Boolean Network-Based Identification of CDX2-Linked Differentiation Targets. A**. Schematic of epithelial differentiation in the colonic crypt, where stem cells reside at the crypt base, and fully differentiated goblet cells and enterocytes localize toward the top. **B**. Illustration of the continuum of crypt differentiation, highlighting various progenitor cell states along the axis from stemness to terminal differentiation. **C**. Overview of the Clustered Boolean Implication Network (BoNE) workflow. Diverse transcriptomic datasets are integrated to generate clusters of co-expressed genes, which are interconnected through one of six invariant Boolean relationships. **D**. Boolean network centered on *CDX2* as a seed gene, generated using the computational platform, Boolean Network Explorer (BoNE[1]), using a pooled microarray dataset (1662 CRCs, 68 adenomas, 170 normal colon tissues). Directed edges indicate Boolean relationships. **E-F**. Machine learning-driven target prioritization. Boolean clusters directly connected to *CDX2* (clusters #2 and #3) were evaluated by linear regression on test datasets. Top five genes (E) from each cluster were selected based on their ability to distinguish CRC from normal tissue (F: GSE dataset identifiers, bar plots, and ROC-AUC values shown). **G**. ROC-AUC analysis showing the ability of Boolean clusters #2 and #3 (which include *PRKAB1* and *CDX2*) to distinguish normal from transformed (and/or inflamed colitic) states in murine colitis models based on gene expression patterns.

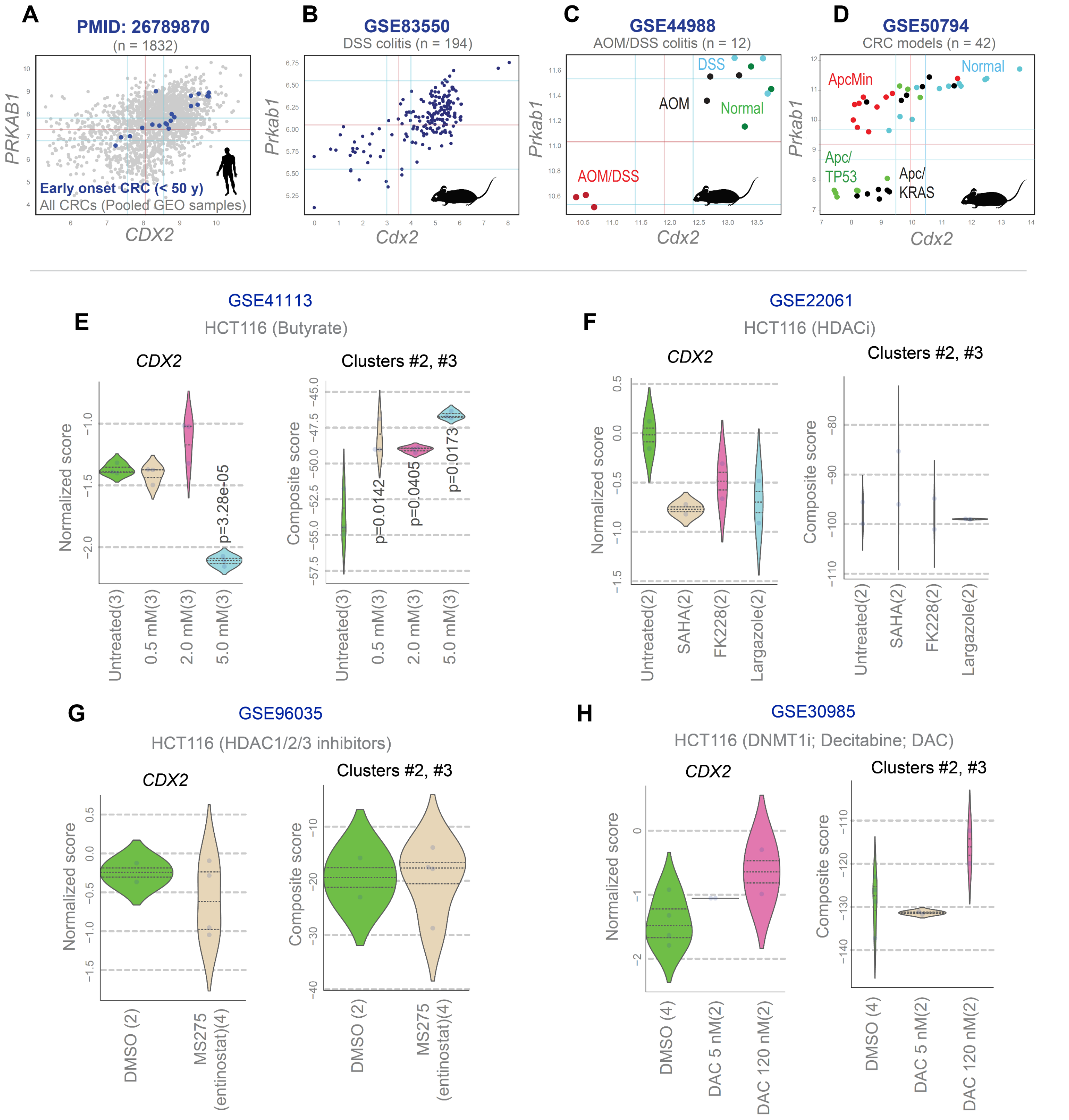


**Figure S2** [Related to Figure 1].

**PRKAB1 Exhibits a Conserved Universal Relationship with and Predicted to Restore CDX2 when Prior CDX2-Restorative Therapeutics have Failed. A-D**. Scatter plots demonstrating a universal equivalent Boolean relationship between *PRKAB1* and *CDX2* across multiple pooled transcriptomic datasets from human and murine models of colorectal cancer (CRC) and colitis. **E-H**. Violin plots show the expression pattern of *CDX2* (left) and network-derived gene clusters #2 and #3 (right) as an effect of various treatments in HCT116 cell lines. *Statistics*: p-values were calculated by Welch's t-test (compared to untreated or control sample#1 in each plot). Only significant p-values are shown.

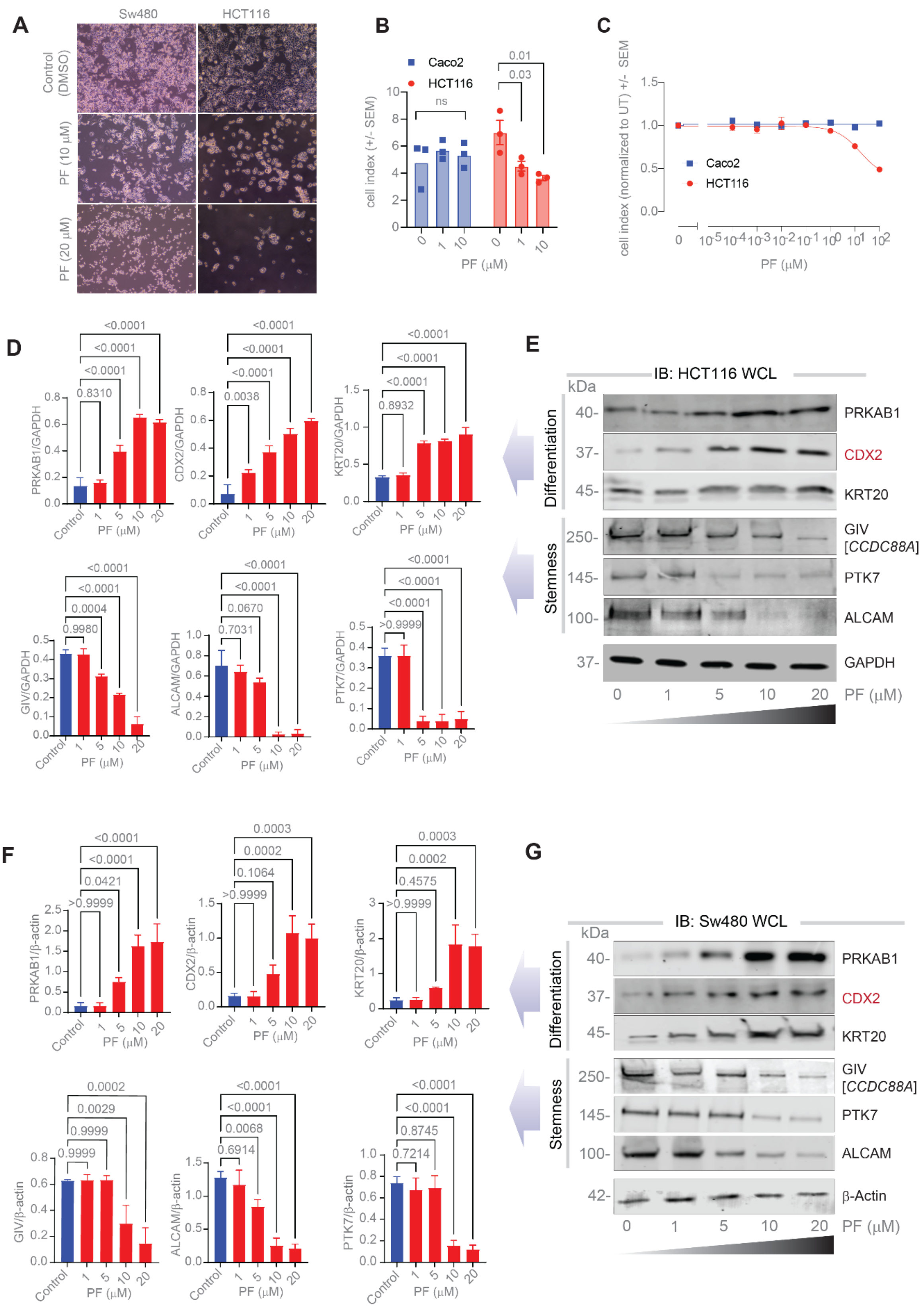


**Figure S3** [Related to Figure 2].

**PRKAB1 Agonist (PF) Induces Differentiation in CRC Cell Lines. A**. Bright-field images (10×) of CRC cell lines treated with 10 μM and 20 μM PF compound for 16 hours. **B-C**. Bar plot (B) showing cell index, measured as impedance, for CDX2-high Caco2 cells and CDX2-low HCT116 cells after 128 h of incubation with varying concentrations of PF compound. Line plot showing normalized cell index (relative to untreated control) during 128 h of incubation with varying concentrations of PF compound. **D-G**. Immunoblots and quantification of HCT116 WCLs (D-E) and Sw480 WCLs (F-G) treated with increasing PF doses for 16 hours prior to lysis (when there is <1-2% cell death). *Statistics*: p-values were calculated using one-way ANOVA. Error bars represent S.E.M. from three independent biological replicates.

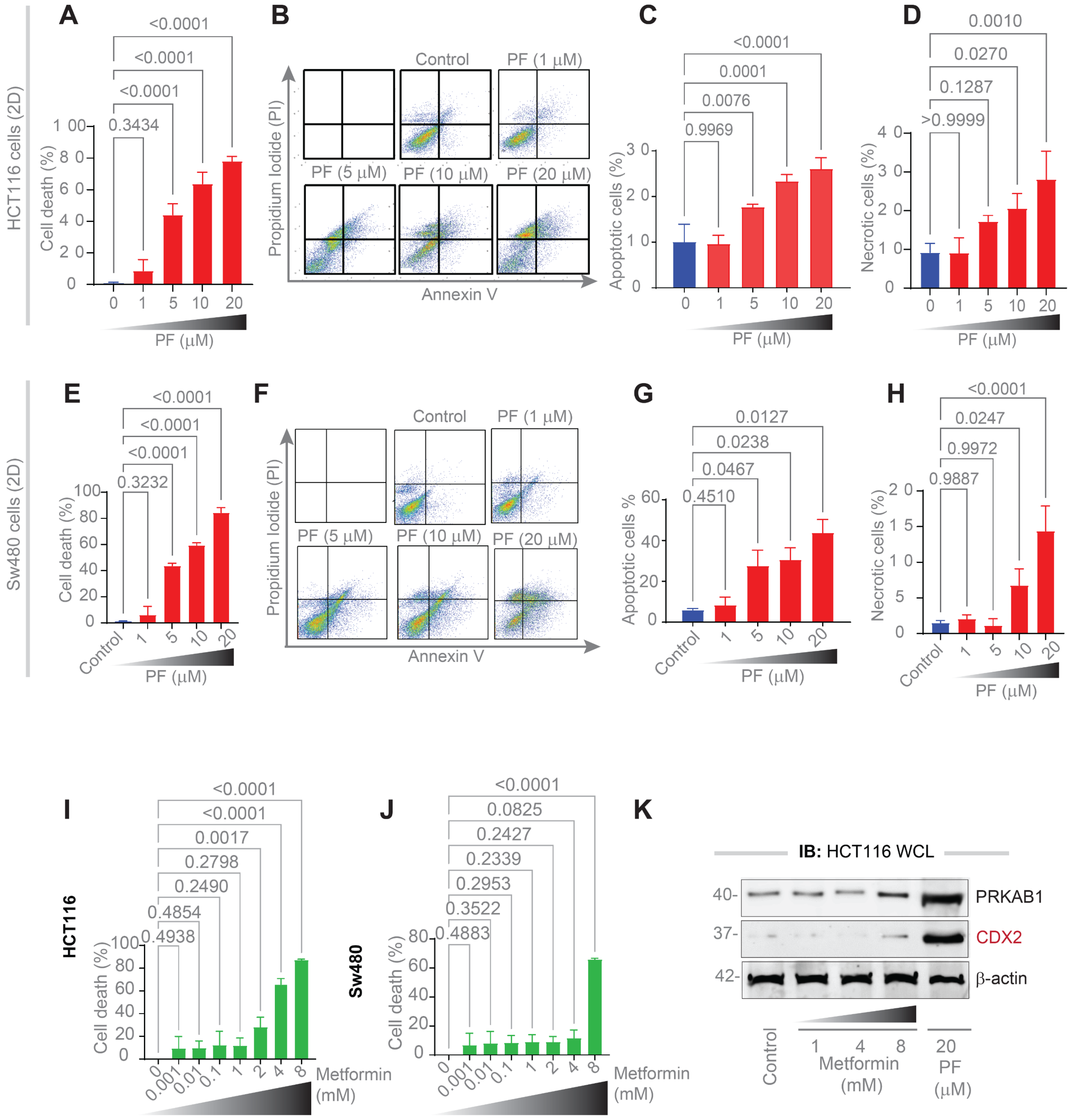


**Figure S4** [Related to Figure 2].

**β1-AMPK Specific Agonist (PRKAB1-agonist, PF), but Not the Indirect AMPK Activator Metformin, Induces Dose-Dependent Cell Death. A-H.** Monolayers of HCT116 (A-D) and Sw480 (E-H) cells were treated with increasing concentrations of the PRKAB1 agonist PF and assessed for apoptosis using Annexin V staining followed by flow cytometry. Representative scatter plots from flow cytometry are shown in B and F. **I-J**. Monolayers of HCT116 (I) and Sw480 (J) cells were treated with various concentrations of the indirect AMPK activator Metformin, and cell viability was measured via MTT assay. **K**. Immunoblots of equal aliquots of whole cell lysates (WCL) of HCT116 cells treated with increasing doses of Metformin and 20 μM PF for 16 h prior to lysis (when there is virtually no cell death). β-actin was used as loading control. *Statistics*: p-values were calculated by one-way ANOVA. Error bars indicate S.E.M of 3 independent biological replicates.

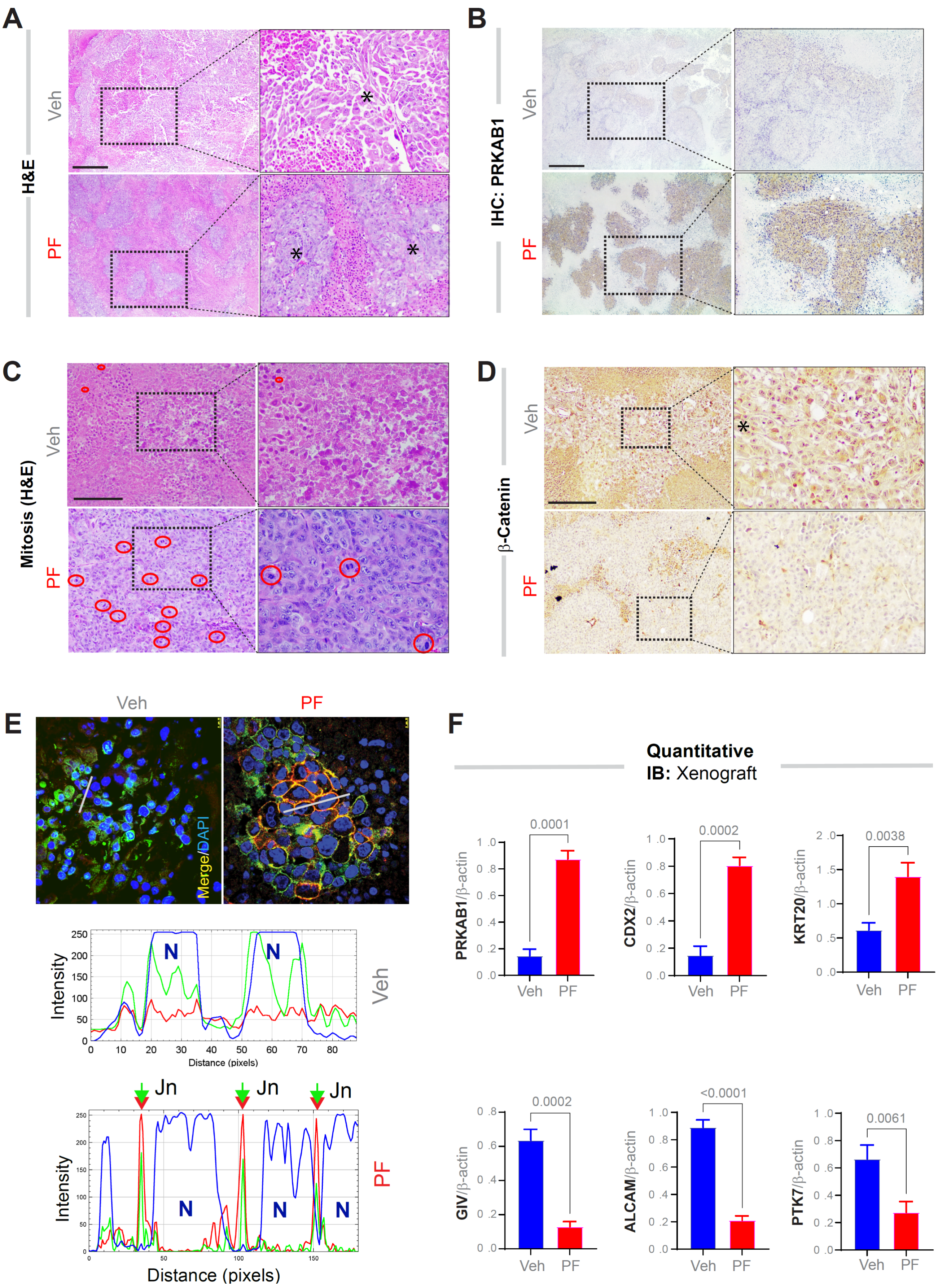


**Figure S5** [Related to Figure 2].

**PRKAB1-agonist (PF) Promotes Differentiation in Xenotransplants. A.** Representative H&E-stained sections of FFPE tumor xenografts. Vehicle (Veh) treated tumors exhibit poorly differentiated, rhomboid-shaped cells arranged in sheet-like patterns with minimal extracellular matrix. In contrast, PF-treated tumors display focal regions of glandular architecture with prominent cell-cell contacts. Asterisks (*) denote viable tumor cell islands; red circles in panel C highlight mitotic figures. **B-D**. Representative fields of H&E and IHC-stained FFPE tumor sections. Scale Bars = 100 µm. **E**. Representative merged images (top) of xenograft sections co-stained for β-catenin (green), E-cadherin (red), and DAPI (blue). RGB intensity plots (bottom, generated in ImageJ) demonstrate co-localization of β-catenin and E-cadherin at cell-cell junctions, indicating membrane-localized β-catenin specifically in PF-treated tumors. Scale bar = 10 µm. See individual channels in **Figure 2N**. **F**. Quantification of differentiation and stemness markers from immunoblots of xenograft lysates in **Figure 2O**. *Statistics*: p-values were calculated by paired t-test. Error bars indicate S.E.M of 3 independent biological replicates.

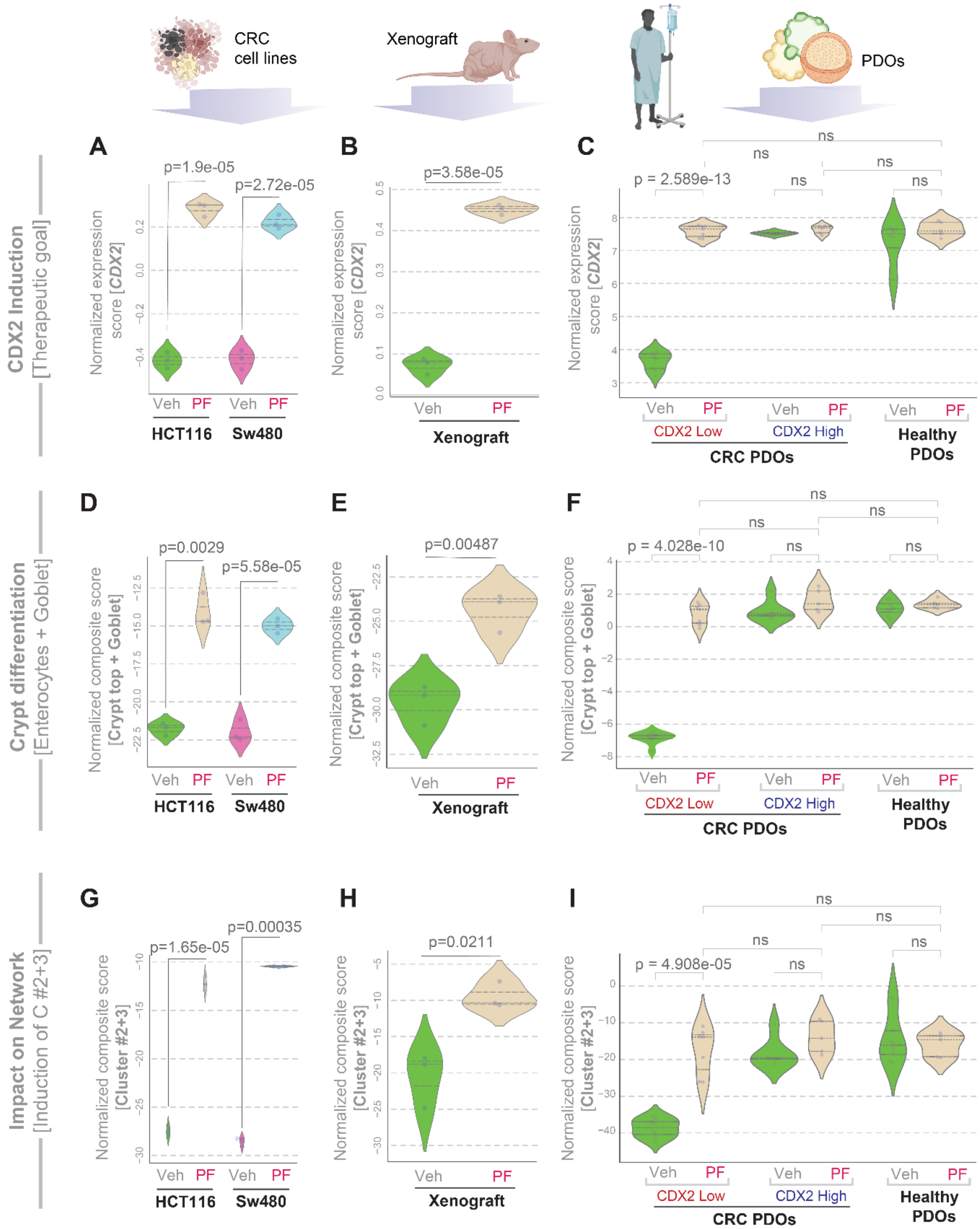


**Figure S6 [Related to Figure 4].**

**PF treatment induces *CDX2*, markers of crypt differentiation, and genes in cluster #2 + #3 in the CRC network in CRC cell lines, xenografts, and PDOs.** Violin plots display the normalized expression of *CDX2* (A-C), markers of absorptive enterocytes and Goblet cells indicative of crypt differentiation (D-F), and composite score of genes in clusters #2 and #3 from proposed CRC network (G-I) in CRC cell lines (A, D, G), tumor xenografts (B, E, H) and PDOs (C, F, I). See **Supplemental Information 2** for a compete catalogue of genes in clusters #2 and #3 and the markers of crypt differentiation. *Statistics*: p-values were calculated by Welch's t-test.

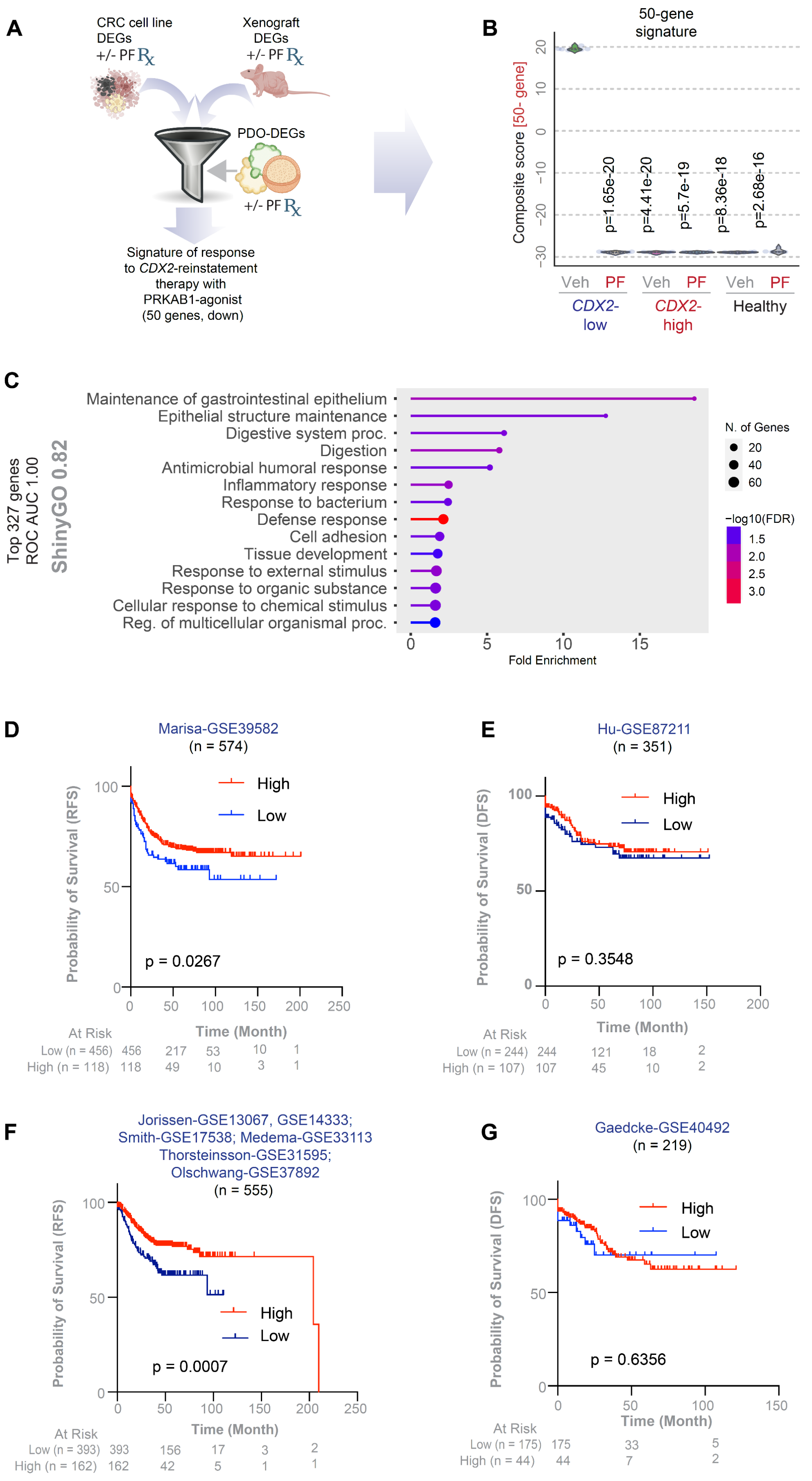


**Figure S7 [Related to Figure 5].**
**A gene signature of therapeutic response predicts better survival in patients.**

**A.** Computational workflow used in the derivation of a 50-gene signature of therapeutic response using an integrated DEA approach. Differentially expressed genes (DEGs) from CRC cell lines and xenografts (see **Supplemental Information 4-5**) were further refined based on their ability to accurately classify PDO samples with/without PF treatment (see **Supplemental Information 6** for gene list rank-ordered by ROC-AUC score). **B.** Violin plots show composite scores of the 50-gene signature in PDOs. p-values are based on Welch's t-test compared to vehicle (Veh)-treated *CDX2*-low PDOs. **C**. Lollipop plots depict pathway enrichment for the top 327 differentially expressed genes (DEGs) induced upon PF treatment in CRC cell lines and xenografts, which also achieved 100% classification accuracy (i.e., ROC AUC 1.00; **Supplemental Information 6**) in PDO samples. **D-G.** Kaplan-Meier plots showing probability of survival (DFS/RFS) over time in various cohort, segregated by high vs low expression scores of *CDX2*, computed using the *StepMiner* algorithm[2] within each cohort. p-value was determined by Log Rank test. See **Supplemental Information 7** for the corresponding O.R.

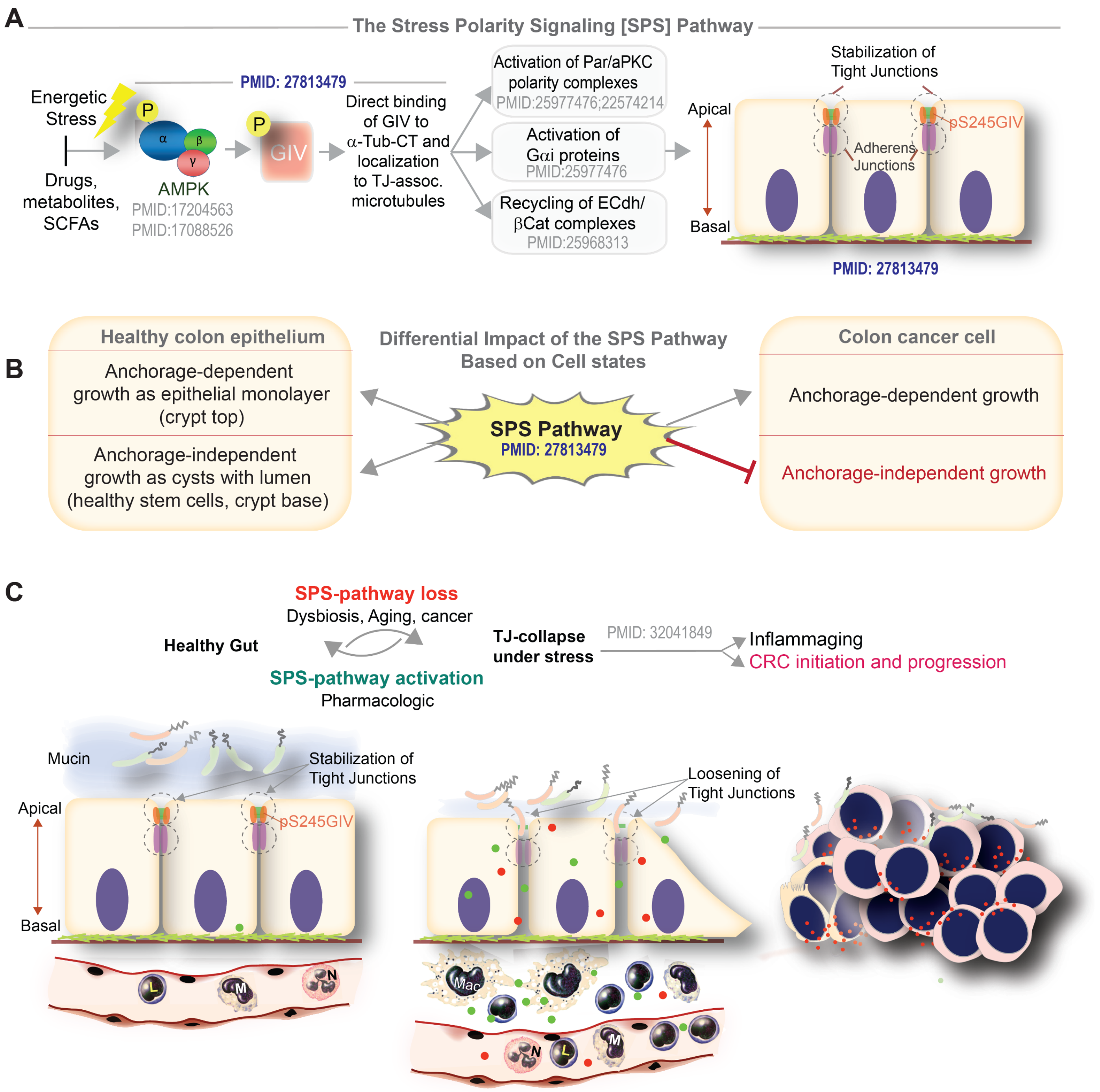


**Figure S8** [Related to Figure 6].

**The Stress Polarity Signaling (SPS) Pathway Maintains Epithelial Polarity in Health and Restrains Malignant Transformation.**

**A.** Schematic overview of the SPS pathway[3-5]. Activation of a junction-localized β1-specific pool of trimeric αβγ-AMPK leads to phosphorylation of GIV at a Serine in position 245 (pSer$^{245}$), which in turn activates Par/aPKC complexes and Gαi proteins. This cascade promotes the recycling of E-cadherin/β-catenin complexes, thereby stabilizing tight junctions and reinforcing apical polarity in epithelial cells. Corresponding PMID#s are listed.

**B.** Diagram illustrating previously published[6] work on the cell state-dependent cell fates induced by the SPS pathway. In healthy epithelial cells, SPS pathway activation permits anchorage-independent cystic growth with preserved lumens and anchorage-dependent growth as monolayers with enhanced barrier function. By contrast, activation of the SPS pathway in SPS-deficient transformed CRC cells is permissive only to anchorage-dependent growth as polarized monolayer but virtually abolishes growth in anchorage independent mode.

**C.** Schematic summarizes published work on the status of the SPS pathway in health and disease. In healthy intestinal epithelium, physiological or drug-induced SPS activation enhances apico-basal polarity and tight junction integrity via GIV phosphorylation. In pathological states (e.g., dysbiosis, aging, or cancer), SPS pathway is suppressed or lost, leading to tight junction breakdown, compromised barrier function, and increased susceptibility to inflammation and CRC progression.
